# Scaling up orphan crop research: A global genetic perspective of cowpea (*Vigna unguiculata*) diversity from 10,617 accessions

**DOI:** 10.1101/2025.11.03.686174

**Authors:** Sofie M. Pearson, Adrian Hathorn, Shichao Sun, Alan Cruickshank, Tracey Shatte, Paulino Munisse, Mercy Macharia Wairimu, Joann Conner, Anna M.G. Koltunow, Jean-Philippe Vielle-Calzada, Peggy Ozias-Akins, Takayoshi Ishii, Matteo Dell Acqua, Sally Norton, Yongfu Tao, David Jordan, Emma Mace

## Abstract

Cowpea (*Vigna unguiculata* L. Walp) is a dryland legume crop, providing essential food and nutritional security for millions of people across the semi-arid tropics, in Africa, Asia, and Latin America. However, as a typical “orphan crop”, cowpea has long remained underrepresented in global genomic research to support crop improvement. Here, we conducted the largest genetic diversity analysis of cowpea to date, comprising 10,617 accessions sourced from seven international collections. Using genotyping-by-sequencing, we characterized the global patterns of genetic diversity, assessed redundancy within and across collections, and examined the geographic structure of the cowpea global allele pool. Our results revealed nine distinct genetic groups with clear geographic associations and fine-scale population differentiation, reflecting dispersal history, regional adaptation and the influence of modern breeding. Duplication across collections was detected, highlighting the need for improved curation and integration of germplasm resources. Landraces from sub-Saharan Africa do not fully capture the genetic diversity present in several other geographic regions, indicating the existence of abundant and untapped genetic resources worldwide. These findings not only provide insights into the genetic structure and evolutionary history of cowpea but also offer a valuable foundation for harnessing global germplasm diversity to enhance breeding potential and accelerate crop improvement.

## INTRODUCTION

Global food security remains a pressing issue, with an estimated one billion people currently underfed, a situation that is likely to intensify as the global population is predicted to surpass nine billion by 2050 (Gu et al., 2021). Meeting this future demand will require increasing food production while meeting sustainability objectives (Tilman et al., 2011, van Dijk et al., 2021). However, relying solely on improvements in major staple crops is unlikely to meet this target (Ray et al., 2013). Diversifying food sources is therefore imperative to ensure nutritional security (Gruber, 2017). Cultivating orphan crops (those which lack advanced scientific research and breeding) offers a promising avenue for diversification and sustainability. Orphan crops include cereals, legumes and root crops often cultivated in limited areas in Africa, Asia and Latin America, play an important cultural role and are vital for the nutrition and livelihood of many households (Ye et al., 2021). These crops possess desirable traits such as resilience to extreme climatic conditions and high nutritional value (Tadele, 2019). Additionally, plant genetic resources including landraces and wild crop relatives conserved in international genebanks are a valuable reservoir of genetic diversity, offering access to novel allelic variation that could contribute in sustainable intensification of yields, providing resistance to diseases and abiotic stress, and increasing nutritional quality (Ye et al., 2021).

Genebanks play a critical role in safeguarding plant genetic resources by storing seeds from diverse species, including orphan crops. These large diverse collections promote conservation and support plant breeding and crop improvement, especially in the face of changing environmental conditions (Asdal et al., 2018). This includes maintaining collections of crop seeds, including traditional and heirloom varieties that may possess traits such as pest resistance, drought tolerance, and adaptability to low-input environments (Hay et al., 2013). Effective management of these genetic reservoirs is crucial for developing cultivars capable of addressing emerging agricultural challenges (Tanksley et al., 1997). Therefore, accurate identification and conservation of genetic diversity within collections are essential for optimizing breeding programs (Mascher et al., 2019). Previous studies assessing diversity in genebanks include major staple crops such as wheat (Schulthess et al., 2022), barley (Milner et al., 2019) and rice (McCouch et al., 2012), but limited studies have assessed the diversity in orphan crops.

The largest genebank collections of cowpea (*Vigna unguiculata* L. Walp) include the International Institute of Tropical Agriculture (IITA) in Nigeria, the United States Department of Agriculture (USDA) National Plant Germplasm System (NPGS) in the United States and the National Bureau of Plant Genetic Resources (NBPGR) in India. Cowpea is a globally important grain legume crop and contributes to food and nutritional security for hundreds of millions of households in the semi-arid tropics, including Africa, Asia and Latin America. Cowpea is an orphan crop and is cultivated mainly for its grains which provide a main source of protein to hundreds of millions of people in developing countries (Boukar et al., 2019). Cultivated cowpea are grouped under *V. unguiculata* ssp. *unguiculata*, which is subdivided into five cultivar groups (c.g.) based on seed and pod traits: (1) *Unguiculata* (cowpea or black-eyed bean, widely cultivated as a pulse or vegetable with more than 16 ovules per pod); (2) *Biflora/Cylindrica* (catjang, common in India, primarily used for forage and has short erect pods with less than 17 ovules per pod and smooth seed); (3) *Sesquipedalis* (yardlong or asparagus bean, common in Asia, grown as a vegetable with very long pods); (4) *Textilis* (a rare form once cultivated for fibre in Africa with long floral peduncles); and (5) *Melanophthalmus* (grown mostly in the Americas, has less than 17 ovules per pod and includes blackeye pea types) (Pasquet, 2000). Cowpea seeds, leaves and green pods are all utilised and provide essential micronutrients such as zinc and iron, which are often lacking in other legumes (Jayathilake et al., 2018, Abebe et al., 2022).

In sub-Saharan Africa, smallholder farmers are the main producers and consumers of cowpea (Bolarinwa et al., 2022) and often grow it as an intercrop with cereals and other crops (Ongom et al., 2023). Cowpea compensates for the nitrogen loss by cereals through sustainable nitrogen fixation in farming systems (Kussie et al., 2024). Cowpea’s centre of domestication is in Africa; from there it has spread to all continents and exhibits high resilience to drought environments and poor soil conditions. However, yields are often adversely affected by cowpea’s susceptibility to pests (such as aphids, weevils and pod borers) and diseases (such as Anthracnose and fusarium wilt). These yield challenges pose a threat to global food security, highlighting the need for continuous breeding efforts. Characterizing global germplasm is critical to explore the untapped resource to meet the challenges in cowpea breeding.

Over the last few decades, significant progress has been made in characterizing the genetic diversity of cowpea. Genetic studies have consistently highlighted West Africa as the primary centre of cowpea diversity, with substantial variation also found across Southern and Eastern Africa, Asia, and the Americas (Fang et al., 2007, Huynh et al., 2013, Xiong et al., 2016, Sodedji et al., 2021, Fiscus et al., 2024). Multiple studies focused on specific geographic regions including Western Africa (Egbadzor et al., 2014, Gbedevi et al., 2021) Southern Africa (Nkhoma et al., 2020, Chipeta et al., 2025, Macharia et al., 2025), Eastern Africa (Desalegne et al., 2016, Ketema et al., 2020), the Iberian Peninsula (Carvalho et al., 2017), China (Chen et al., 2017) and Korea (Seo et al., 2020). However, the relationship among cowpea accessions from a comprehensive global geographic distribution remains limited. There have been efforts at characterising global core collections (Mahalakshmi et al., 2007, Fatokun et al., 2018, Muñoz-Amatriaín et al., 2021), however the bulk of these collections constitute traditional cultivars from West Africa and have limited sampling from continents outside Africa, leaving gaps in understanding the full extent of global cowpea diversity. There is a need for studies to integrate larger and geographically diverse collections to capture the untapped genetic variation critical for breeding programs, and in positioning cowpea as a valuable genetic resource and transforming it from an underutilised crop into a key contributor to global and nutritional security.

Here we explore a curated selection of the global gene pool of cowpea to understand patterns of genetic diversity and population structure. The collection is particularly valuable because it integrates accessions from five genebanks and two collections, capturing a broad spectrum of geographic origins and genetic backgrounds. We assembled a dataset of 10,617 accessions, approximately 23% of the total global cowpea accessions, representing both core and diverse germplasm from six continents and 121 countries, and applied cost-effective and robust reduced-representation sequencing to generate high-quality genotypic data. The main objectives of this study were to (1) Characterise a global collection of cowpea from seven international collections, (2) Identify identical accessions within and between collections using genotypic data, and (3) Assess the composition of the global cowpea collection in the context of diversity and genetic structure.

## MATERIALS AND METHODS

### Plant Materials

A total of 10,617 cultivated cowpea (*Vigna unguiculata*) accessions and one sister subspecies accession (*Vigna unguiculata* subsp. *stenophylla*) were used in this study. These accessions originated from diverse geographic regions and seven collections. The majority of these accessions were from the United States Department of Agriculture (USDA) Genebank (63%; *n* = 6,735), followed by the International Institute of Tropical Agriculture (IITA) Genebank, Nigeria (18%; *n* = 1,942), representing the core collection of cowpea accessions described by Mahalakshmi et al. (2007). A collection of accessions held at Australian Grains Genebank (AGG), Australia (7%, *n* = 791), and the National Agriculture and Food Research Organization (NARO) Genebank, Japan (4%, *n* = 414), were included to expand geographic distribution. The diversity of the cowpea accessions was further enriched by the inclusion of several smaller but highly valuable collections from various regions. These included 351 accessions from the University of California, Riverside (UCR) mini-core collection (Muñoz-Amatriaín et al., 2021), 271 accessions from the National Genebank of Mozambique at the Instituto de Investigação Agrária de Moçambique (IIAM) (Macharia et al., 2025), and 114 accessions from the Central and Southern American collection from Unidad de Genómica Avanzada, Langebio Cinvestav, Mexico (LAN). Together, these diverse sources represent a broad spectrum of genetic variation, including both regional and global cowpea germplasm, thereby providing a comprehensive foundation for subsequent genetic studies. Passport descriptors (i.e., metadata) including taxonomic classification, location of provenance, collection site and improvement status were collated for all accessions from their respective collection, where available. Metadata were accessed online through: the USDA-ARS Germplasm Resources Information Network (GRIN) (https://www.ars-grin.gov/), IITA Genebank (https://my.iita.org/accession2/browse.jspx), Australian Grains Genebank GRIN-Global Database System (https://ausgenebank.agriculture.vic.gov.au/gringlobal/search), NARO (https://www.gene.affrc.go.jp/databases-plant_search_en.php), Genesys (https://www.genesys-pgr.org/) and supplementary material from Muñoz-Amatriaín et al. (2021) and Macharia et al. (2025). Metadata for accessions used are provided in Table S1.

### Genotyping-by-sequencing

Genotyping of the global cowpea collection was conducted by Diversity Arrays Technology (DArT) Pty Ltd (https://www.diversityarrays.com/services/dartseq/) using the DArTseq™ reduced-representation sequencing platform (Kilian et al., 2012). For each accession, 30-40mg of seed tissue derived from single or multiple seeds was sampled and placed in a 96-well plate. Samples were ground in a TissueLyser II (QIAGEN, Hilden, Germany) using metal rods at 30 Hz for 20-30 seconds, repeated twice. Following grinding, the rods were removed and tubes sealed. Ground samples were stored at 4°C before shipping to DArT where DNA extraction was conducted using a modified cetyl trimethyl ammonium bromide (CTAB) method (Doyle et al., 1987). For accessions restricted by quarantine regulations, leaf material from the NARO accessions were sampled following the procedure described by Ofem et al. (2025) and sent to DArT for DNA extraction. Samples from the IIAM collection were extracted at the Genebank in Mozambique using the protocol described by Macharia et al. (2025) and subsequently shipped to DArT for sequencing. All DNA samples were then digested with methylation sensitive restriction enzymes (*Pst*I and *Mse*I) to reduce genome complexity and to remove repetitive sequences (Kilian et al., 2012). Libraries with size lengths of 200-500bp were constructed using a TruseqNano DNA HT sample preparation kit (Illumina; catalog no. FC-121-4003) following the manufacturer’s recommendations, and sequenced on an Illumina HiSeq 2500 platform to produce 77bp paired end reads (Sansaloni et al., 2011). Raw sequencing reads were processed using DArT’s proprietary analytical pipeline. In this pipeline, the FASTQ files were filtered to remove adapters, low-quality reads, and redundant reads with barcode regions stringently filtered (Phred pass score ≥ 30) to ensure accurate sample assignment to the correct reads. Identical reads were collapsed into ‘fastqcall’ tag files which were then used as input for DArT’s secondary pipeline for SNP analysis (Raman et al., 2014). Reads were subsequently aligned to the cowpea reference genome IT97K-499-35 v1.2 (Lonardi et al., 2019) using BWA-MEM v0.7.17 (Li et al., 2009), and SNP genotypes were called with DArT’s proprietary secondary pipeline (DArTsoft14) (Sansaloni et al., 2020). Genotype data were delivered in VCF format (Cruz et al., 2013). During post-DArT quality control, samples with a call rate ≤ 0.50 and markers with a call rate ≤ 0.80 were excluded. The overall call rate in the raw genotypic dataset was approximately 82%. Missing genotypes were inferred using Beagle v 5.0 (Browning et al., 2016). Further filtering was applied by removing markers with heterozygosity ≥ 0.4 and individuals with heterozygosity ≥ 0.2. The final filtered dataset consisted of 10,618 accessions and 4,290 high-quality SNPs, denoted the “raw dataset”, providing a robust dataset for subsequent analyses of genetic diversity. The genomic distribution and density of these SNPs were visualised using the online SNP Density tool (https://www.bioinformatics.com.cn/en) (Figure S1).

### Genetic relatedness between collections and dataset generation

Cultivated cowpea were extracted from the raw dataset and markers with a minor allele frequency ≥ 0.01 were retained using VCFtools v 0.1.16 (Danecek et al., 2011), resulting in 3,073 SNPs across 10,617 accessions denoted the “cultivated dataset”. An initial principal component analysis (PCA) was performed using the *prcomp* function from the stats package in R v 4.3.0 (R Core Team, 2023) to visualize accession distribution according to their metadata, including cultivar group, improvement status, collection (i.e. germplasm source) and geographic provenance.

Duplicate accessions within the cultivated dataset were identified based on three metrics: genetic distance, genomic relatedness and identity by descent (IBD). Pairwise Nei’s genetic distance was calculated using the *nei.dist* function from the poppr package v 2.9.4 (Kamvar et al., 2014), ranging from 0 (minimal genetic difference) to 1 (maximal genetic difference) using the cultivated dataset. A conservative threshold of ≤ 0.03 was used to identify genetically similar pairs. Van Raden’s genomic relationship matrix was calculated using the *Gmatrix* function from the AGHmatrix package v 2.1.4 (Amadeu et al., 2023) using the cultivated dataset, with pairwise values ≥ 1.3 considered indicative of genetic relatedness. Linkage disequilibrium (LD) pruning was performed using PLINK v 1.90b7.2 (Purcell et al., 2007) on the cultivated dataset with the parameter *--indep-pairwise 50-5-0.5* to generate the “cultivated-pruned dataset”, consisting of 2,302 unlinked SNPs. This dataset was used to calculate identify by state (IBS), defined as the average proportion of alleles shared between pairs of individuals at genotyped SNPs. The proportion of whole genome IBD was also calculated using the *--genome* flag and PLINK’s Hidden Markov Models (HMM) algorithm (Purcell et al., 2007). Accessions with a *π̂* value greater than 0.5 indicated parent/child relationships, sib-pairs and duplicated accessions. Accession pairs would need to fit all three criteria to be considered a potential duplicate: genetic distance ≥ 0.03, genomic relatedness ≥ 1.3 and *π̂* ≥ 0.6. To adopt a more conservative approach, passport metadata were also considered: accession pairs that met all three genetic criteria were required to share the same accession name or alias. Potentially duplicate accessions with shared names were thus identified as duplicates and removed from the raw dataset using bcftools v 1.15.1 (Danecek et al., 2021). This dataset was filtered to remove markers with a minor allele frequency ≤ 0.01 resulting in 3,083 SNPs and 9,609 accessions and was denoted the “non-redundant dataset”. The non-redundant dataset was also LD pruned using the methods described above, resulting in 2,312 SNP retained to generate the “non-redundant pruned dataset”.

In the cultivated-pruned dataset, relatedness was summarised as the proportion of accession pairs with *π̂* > 0 within and between collections, reported as ‘percent pairwise IBD’ in Figure 1b. Relatedness was calculated for all pairs between 10,617 accessions using PLINK v 1.90b7.2 (Purcell et al., 2007) by summing the pairs with *π̂* > 0 within a collection combination, then calculating the total number of pairs within the collection combination and finally calculating the percentage of related pairs relative to the total possible combinations for the collection combination. In addition, the distribution of *π̂* values within each geographic provenance were assessed with density plots in R, reported as ‘pairwise IBD values’ in Figure 2a and 2b.

**Figure 1.**
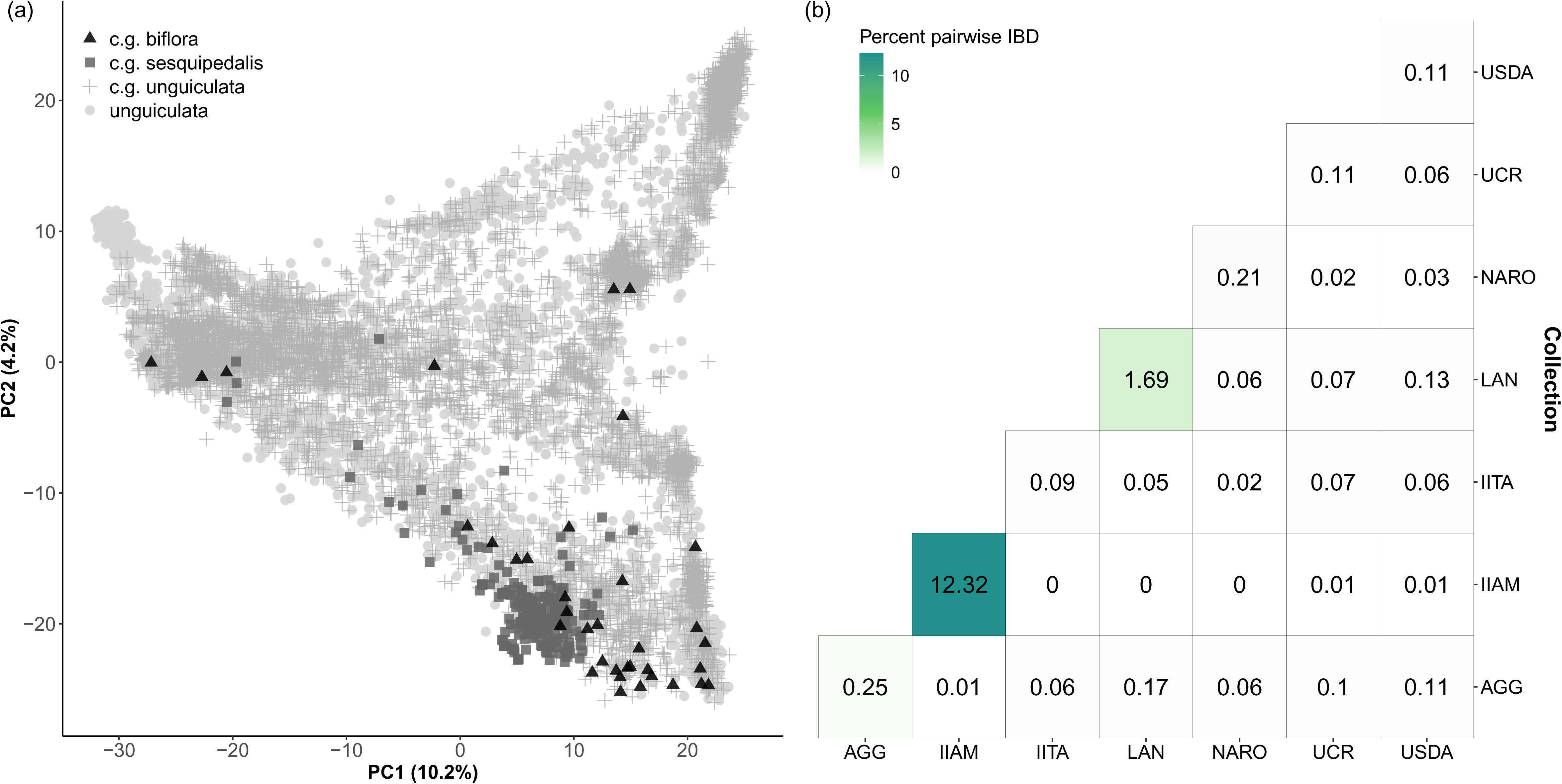
Genetic diversity of the cultivated global cowpea collection. (a) Principal Component (PC) Analysis of 10,617 accessions color-coded based on cultivar group (c.g.), where black triangles = biflora (*n* = 36), grey squares = sesquipedalis (*n* = 240), grey plus = unguiculata (*n* = 5,752) and grey circles = species level *Vigna unguiculata* or subspecies *unguiculata* combined (*n* = 4,589). (b) Percent pairwise identity by descent (IBD) greater than zero between collections. The percentage is calculated from the total number of pairs between each comparison compared to observed pairwise IBD values greater than zero.

**Figure 2.**
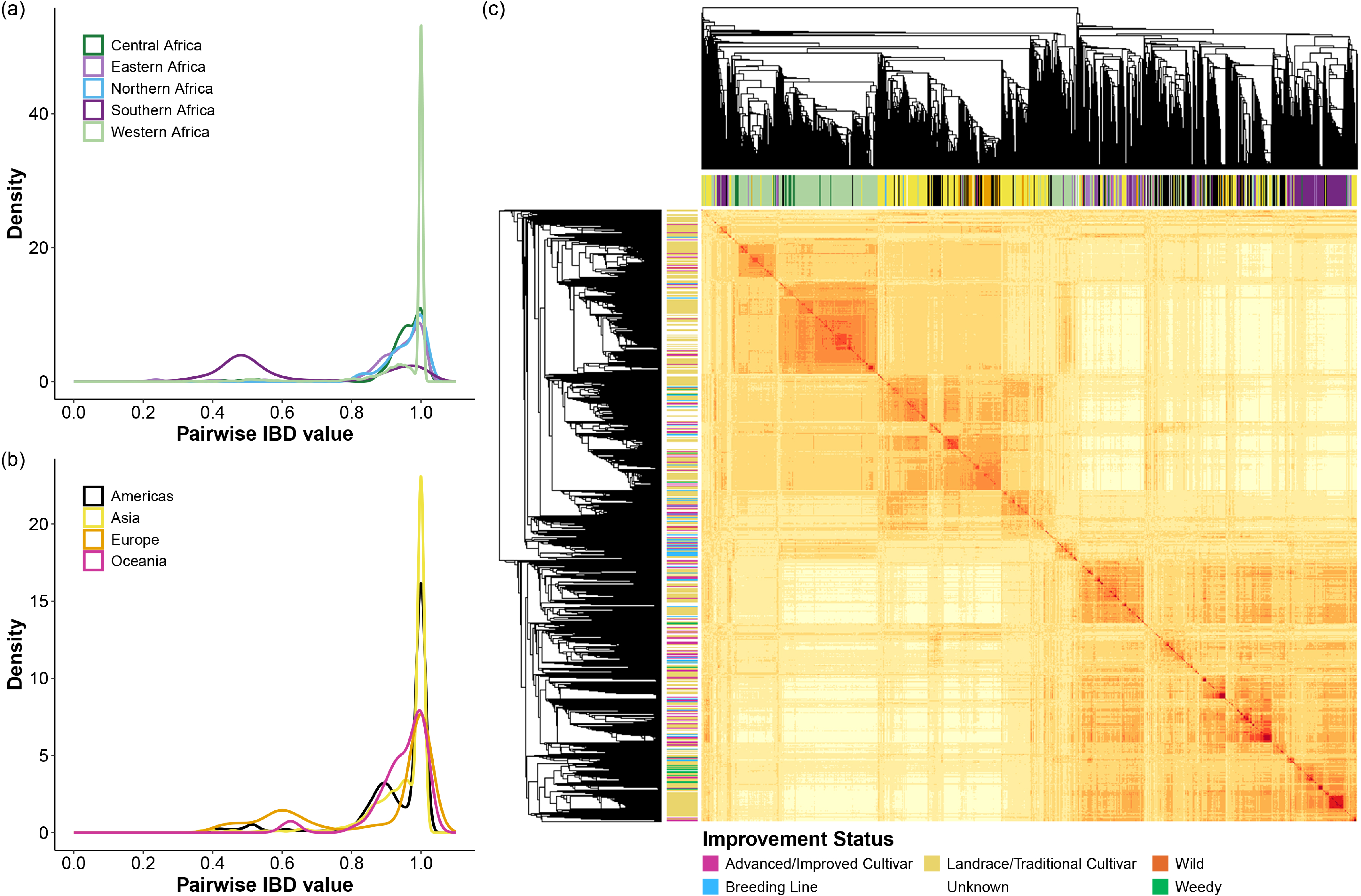
Genetic relatedness of the cultivated global cowpea collection. (a) and (b) Pairwise identity by descent (IBD) frequency distribution for nine geographic regions. (c) Genomic relationship matrix of 10,617 accessions, where yellow = dissimilar and red = similar. Accessions are coloured based on their geographic origin in the columns (color key in (a) and (b)) and improvement status in the rows (color key presented below the heatmap).

### Genetic structure and diversity analysis

To explore the overall patterns of genetic clustering in cowpea germplasm, two separate analyses were performed: (1) Model-based structure analyses used ADMIXTURE v 1.3.0 (Alexander et al., 2009), STRUCTURE v 2.3.4 (Pritchard et al., 2000), and the sparse nonnegative matrix factorization (sNMF) algorithm (Frichot et al., 2014) implemented in the LEA package v 3.12.2 (Frichot et al., 2015); and (2) Phylogeny reconstruction used IQ-TREE v 2.2.2.3 (Minh et al., 2020). The “non-redundant pruned dataset” described above was used for structure analyses and phylogenetic analyses.

ADMIXTURE, STRUCTURE and sNMF were used to perform model-based clustering analyses to explore the broad patterns of population structure and determine the number of putative ancestral populations (*K*). STRUCTURE uses a Bayesian clustering approach, ADMIXTURE uses maximum likelihood modelling, and sNMF uses sparse nonnegative matrix factorization algorithms to estimate individual ancestry coefficients. As ADMIXTURE is computationally faster to run compared to STRUCTURE, the ADMIXTURE analysis was performed using 10 independent runs specified with a different seed using -s with each run set from 1 to 40 with 1-cross-validations (--cv = 10) to determine the *K* values with the lowest cross-validation error. As the cross-validation error continued to decline as *K* increased, the optimal *K* value was determined using the delta *K* method (Evanno et al., 2005). Subsequently, ten independent runs for each simulated value of *K* ranging from 1 to 15 was performed using STRUCTURE. Each run consisted of a burn-in period of 50,000 iterations, followed by 100,000 Markov Chain Monte Carlo (MCMC) replications. The putative optimal *K* was determined using the pophelperShiny app v 2.1.1 (Francis, 2017) using the delta *K* method (Evanno et al., 2005). In parallel, Model-based estimation of ancestral coefficients was performed using the sNMF algorithm (Frichot et al., 2014) implemented in the R package LEA (Frichot et al., 2015). The number of ancestral populations (*K*) was set from 1 to 25 with 20 iterations for each *K* value. The best-fit *K* value was determined by assessing the change in cross-entropy for each *K*, i.e. the elbow of the curve. Ancestral proportions from the sNMF analysis were plotted using the PophelperShiny app v 2.1.1 (Francis, 2017). Accessions were assigned to populations if their individual admixture coefficient matrix (Q) was at least 70%, and those with less than 70% membership probability were considered mixed. Population structure was further examined using the *prcomp* function to perform a PCA. Distribution of genetic groups to country of provenance was visualised in R using the rworldmap v 1.3-8 (South, 2011) and ggplot2 v 3.5.1 packages (Wickham, 2016).

For the phylogeny reconstruction, one *Vigna unguiculata* subsp. *stenophylla* accession was included as the outgroup in the non-redundant pruned dataset. A maximum likelihood phylogeny was generated using IQ-TREE v 2.2.2.3 (Minh et al., 2020) specifying the polymorphisms-aware phylogenetic model (Schrempf et al., 2016, Schrempf et al., 2019). ModelFinder (Kalyaanamoorthy et al., 2017) was first used to determine the best-fit substitution model for the data (GTR+F+ASC+R10). Default parameters were used for tree construction with the *stenophylla* accession assigned as the outgroup. Convergence was not assessed by undertaking independent runs of IQ-TREE and examining tree topology due to the large number of tips (9,610) and nodes, but did ensure that the log-likelihood values were stable at the end of the run. Branch support was performed using 10,000 replicates of UFboot (Hoang et al., 2018). The 9,610 taxa tree was plotted in R using ggtree v 3.8.2 (Yu et al., 2017) and ggtreeExtra v 1.10.0 (Xu et al., 2021).

Relatedness in terms of the proportion of IBD was calculated for all pairwise accessions using the “non-redundant pruned dataset” and the distribution of these proportions were assessed within each group using a density plot in R. To assess genetic diversity, the non-redundant dataset was used to estimate six diversity metrics. Nucleotide diversity (π, (Hohenlohe et al., 2010)), *F*_IS_ (Weir et al., 1984), pairwise *F*_ST_ (Weir et al., 1984), expected and observed heterozygosity (*H*_E_ and *H*_O_, respectively) were calculated for each of the genetic groups and 8 geographic regions using snpR v 1.2.9.2 (Hemstrom et al., 2023). Allelic richness (Hurlbert, 1971) and the number of private alleles for groups and regions were calculated using hierfstat v 0.5-11 (Goudet, 2005), and poppr v 2.9.4 (Kamvar et al., 2014, Kamvar et al., 2015), respectively. AMOVA was used to partition the total variance in allele frequencies within and among hierarchies to assess the significance level at each partition: *K* = 2, *K* = 3, *K* = 9 and collection, and was implemented in the packages poppr v2.9.4 (Kamvar et al., 2014, Kamvar et al., 2015) and ade4 v1.7-22 (Dray et al., 2007).

### Geographic enrichment assessment of genetic groups

Collection metadata was used to derive geographic origin for each accession, with a small number from an unknown origin (*n* = 180). Non-parametric Kruskal-Wallis rank sum tests were used to assess geographic enrichment of each genetic group, and pairwise Wilcoxon rank sum tests with a Bonferroni correction were applied to evaluate significant enrichment based on geography. After assessing broad-scale geographic enrichment, fine-scale geographic regions were assessed using the same two tests using the *kruskal.test* and *pairwise.wilcox.test* from the stats package in R 4.3.0 (R Core Team, 2023). Regions with fewer than 10 accessions, including the British Isles, Micronesia, Polynesia and Western Europe, were removed prior to analysis.

### Code availability

Scripts for genetic relatedness, structure, diversity and geographic enrichment analyses are available in the following GitHub repository: https://github.com/SofiePearson/CowpeaDiversity

## RESULTS

### Metadata-based summary of the global cowpea collection (Taxonomy and Improvement Status)

A total of 10,617 cultivated cowpea (*Vigna unguiculata*) accessions and one sister subspecies accession (*Vigna unguiculata* subsp. *stenophylla*) were used in this study. These accessions originated from diverse geographic regions and seven collections. The majority of these accessions were from the United States Department of Agriculture (USDA) Genebank (63.4%; *n* = 6,735), followed by the International Institute of Tropical Agriculture (IITA) Genebank, Nigeria (18.3%; *n* = 1,942), representing the core collection of cowpea accessions described by Mahalakshmi et al. (2007). The Australian Grains Genebank (AGG), Australia (7.4%, *n* = 791); the National Agriculture and Food Research Organization (NARO) Genebank, Japan (3.9%, *n* = 414); a collection from the University of California, Riverside (UCR), USA (3.3%, *n* = 351) representing the mini-core collection described by Muñoz-Amatriaín et al. (2021); a collection from the National Genebank of Mozambique at the Instituto de Investigação Agrária de Moçambique (IIAM; 2.6%, *n* = 271); and a Central and Southern American collection from Unidad de Genómica Avanzada, Langebio Cinvestav, Mexico (LAN; 1.1%, *n* = 114) also constituted the global collection.

The majority of cultivated cowpea assessed in this study belonged to the *unguiculata* cultivar group (*n* = 5,752), with smaller representations from the *sesquipedalis* (*n* = 240) and *biflora* (*n* = 36) groups (Figure 1a). Forty-three percent of the accessions had passport data at the *unguiculata* species (*n* = 3,670) or subspecies (*n* = 919) level (Table S1). Cultivated cowpea accessions were classified into six improvement categories: Advanced/Improved Cultivar, Breeding Line, Landrace/Traditional Cultivar, Weedy, Wild and Unknown (Figure S2). The AGG collection consists primarily of Breeding Lines (74%), with a small percentage of Landrace/Traditional Cultivars (15%) and Unknown accessions (11%). In contrast, the IITA, IIAM, NARO and UCR collections contain a large proportion of Landrace/Traditional Cultivars (86%, 100%, 79% and 74%, respectively). IITA and UCR also contain Breeding Lines (10% and 22%, respectively), while NARO also contains Unknown (12%) and Weedy (6%) accessions. The LAN collection contains mostly Unknown (98%) and Advanced/Improved Cultivars (2%). The USDA Genebank exhibited greater diversity, including Landraces (42%), Unknown (35%), Advanced/Improved Cultivars (19%), and Wild accessions (3%) (Table S2).

### Identification of breeding material from genomic signatures

Pairwise identity by descent (IBD) was calculated for all 10,617 cultivated cowpea and assessed for differences between collections and geographic provenance. Pairwise IBD values ranged from 0 to 0.17% between collections and from 0.09 to 12.3% within collections (Figure 1b). Higher pairwise IBD was observed between AGG-LAN (0.17%), LAN-USDA (0.13%), and AGG-USDA (0.11%), while no pairwise IBD was detected between accessions IIAM-LAN, IIAM-IITA, and IIAM-NARO. The highest pairwise IBD was found within the IIAM Genebank (12.3%), while all other collections had values below 2%.

All geographic regions have an IBD peak around 0.95 and 1, indicating many similar and near identical accessions (Figure 2a and Figure 2b). Southern Africa and the Americas have a larger IBD peak around 0.5 (Figure 2a and Figure 2b), indicating that there are many breeding lines (parent/child and half-sibs) originating from these regions.

The genomic relationship matrix heatmap (Figure 2c) provides a visual summary of related accessions with individuals color-coded by geographic provenance in the columns and improvement status in the rows. As an example, a large group of accessions in the top left corner from Western Africa consisting of Advanced/Improved Cultivars, Breeding Lines, Landraces/Traditional Cultivars and Unknown accessions are similarly related. Another example is a small group of Landrace/Traditional Cultivars from Southern Africa in the bottom right corner are highly related.

### Duplication within and between collections

Potential duplicate accessions among 10,617 cultivated cowpea were identified using Nei’s genetic distance with a threshold of ≤ 0.03, Van Raden’s genomic relatedness with a threshold of ≥ 1.3, and whole genome IBD with a threshold of ≥ 0.6. Each accession was compared to all other accessions in a pairwise manner for each of the metrics, resulting in a total of 112,710,072 pairwise combinations. Reciprocal pairs (e.g., accession1 vs accession2 and accession2 vs accession1) and identical pairs (e.g., accession1 vs accession1) were removed, leaving 56,355,036 unique combinations. Of these, 33,252 pairs met the three criteria and were classified as “potentially duplicated accessions” (Table S3). A higher proportion of potential duplicates were detected within individual collections (57%) than between different collections (43%). Of these 33,252 pairs, 1,329 were confirmed as “duplicated accessions” based on passport alias names (Table S3). A higher proportion of duplicate accessions with matching names were detected between collections (74%) compared to within collections (26%). In cases where multiple pairs of accessions had an identical name, one or multiple accessions were retained depending on the genetic distance, GRM and IBD values. A total of 9,609 non-redundant accessions were retained for subsequent analyses to assess population structure.

### Global genetic structure of cowpea

#### Genetic structure

The genetic population structure of 9,609 cultivated cowpea accessions in the non-redundant pruned dataset, was inferred using three model-based clustering algorithms implemented in ADMIXTURE, STRUCTURE and LEA (Frichot et al., 2015). Cross-validation values from ADMIXTURE declined steadily with increasing *K*, without supporting a clear optimum, indicating the absence of a clearly defined optimal number of genetic groups in the dataset. However, the log-likelihood and Δ*K* analyses indicated that *K* = 2 showed the best fit for both ADMIXTURE (Figure S3) and STRUCTURE (Figure S4). The sNMF cross-entropy analysis also supported two major populations, with further subdivision at higher *K* values (Figure S5). For all *K* values, an accession was assigned a genetic group if membership assignment to a group was greater than 70% (Table S4), while a lower percentage indicated admixture. At *K* = 2, the structure analysis separated the cowpea germplasm into Population 1 and Population 2. The ADMIXTURE, SRUCTURE and sNMF results at *K* = 2 were identical, so admixture proportions from sNMF are reported. The distribution of cowpea accessions between the two groups indicated that Population 2 (green) had the highest percentage of membership (43%) with 4,115 accessions, while Population 1 (purple) had the lowest percentage of membership (35%) with 3,415 accessions. However, the inferred ancestry indicated that 2,079 accessions (22%) were admixed (Figure 3a and Figure 3d, first panel). Support for *K* = 3, indicated further division of the accessions into three populations. Population 1 remained as Population 1 while Population 2 was divided into two Populations: Population 2 and Population 3 (Figure 3b and Figure 3d, second panel). Population 1 (purple) had the highest percentage of membership (34%) with 3,275 accessions, Population 3 (yellow) had an intermediate percentage of membership (20%) with 1,959 accessions, and Population 2 (green) had the lowest percentage of membership (13%) with 1,266 accessions. Splitting the accessions into three populations resulted in an increase in the number of accessions admixed between populations (*n* = 3,109, 32%).

**Figure 3.**
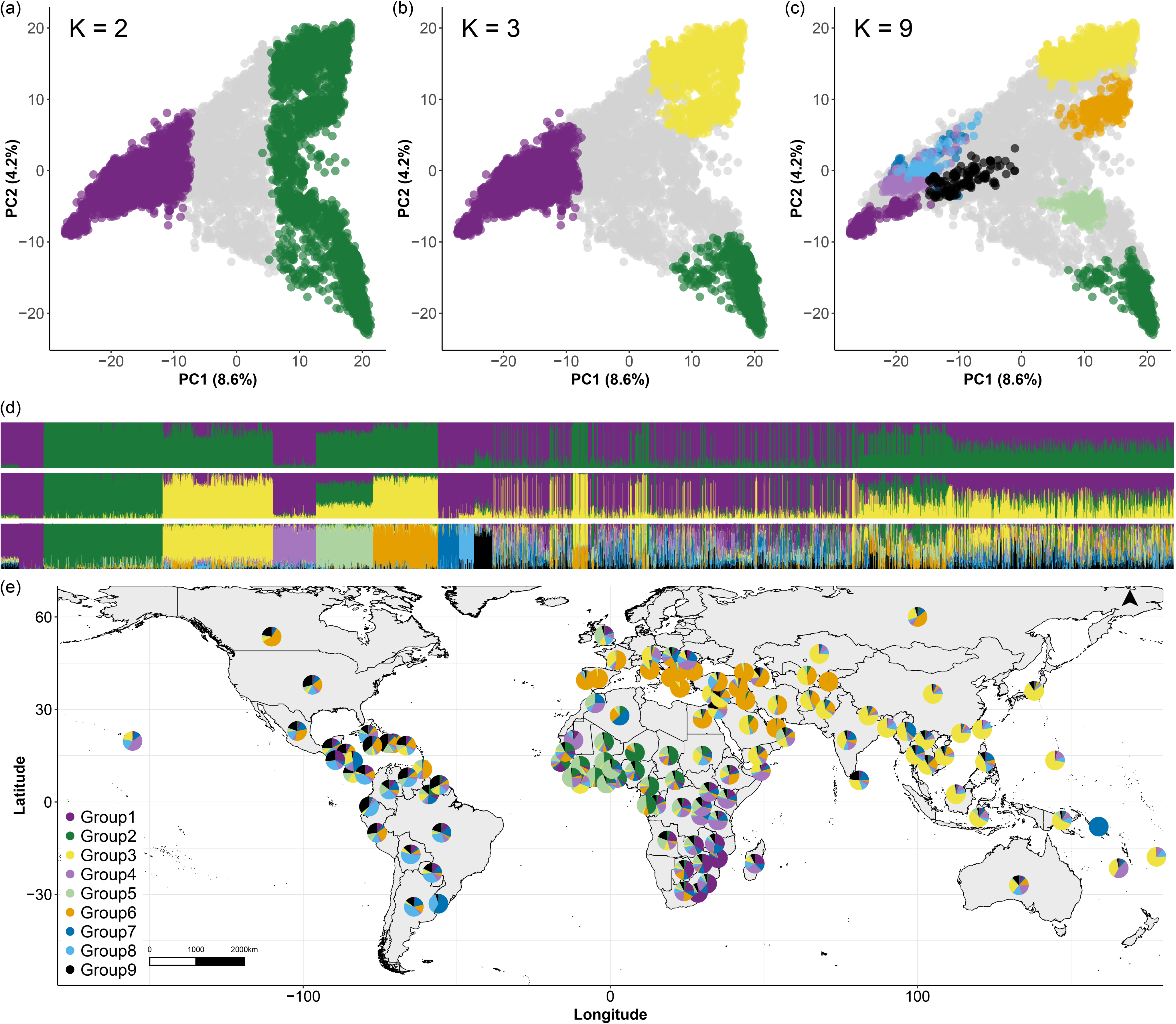
Population structure and geographic distribution of the non-redundant global cowpea collection. (a) – (c) Principal Component (PC) Analysis of the 9,609 accessions coloured by population at *K* = 2, *K* = 3 and *K* = 9. Color key presented in (e). (d) Model-based clustering of *K* = 3, *K* = 3 and *K* = 9. The x-axis has accessions ordered according to their assignment to each *K* = 9 group. The y-axes quantify the membership to each ancestral population represented in colors corresponding to each *K* = 9 group. Color key presented in (e). (e) Geographic distribution of the genetic groups at *K* = 9. For each country, the pie represents the proportion of the ancestral allele frequencies.

The Populations at *K* = 2 and *K* = 3 reflect the broad genetic structuring in cowpea and have general geographic trends. At *K* = 2, accessions belonging to Population 1 originate from Western Africa, Eastern Africa, Southern Africa, and the Americas. Accessions in Population 2 originate from Western and Central Africa, Europe/Mediterranean and Asia (Figure S6). At *K* = 3, when Population 2 was divided into Population 2 and Population 3, a clear distinction emerged, with African accessions in Population 2 and European/Mediterranean and Asian accessions in Population 3 (Figure S7).

We considered additional genetic structure to investigate the potential for population substructure across different regions. Support for a *K* = 9 model further divided the accessions into nine genetic groups (Figure S5). Population 1 split into Groups 1, 4, 7, 8 and 9; Population 2 split into Groups 2 and 5; and Population 3 split into Groups 3 and 6 (Figure 3c and Figure 3d, third panel). Less than half of the 9,609 accessions (*n* = 4,025) were assigned to a group using a 70% membership threshold, with the remaining 5,585 accessions admixed among groups (Figure S8).

#### Geographic enrichment of genetic groups

Geographic enrichment of the nine genetic groups was assessed using non-parametric tests, focusing on group proportions rather than individual assignments. Five groups exhibited significant geographic separation: Group 2 in Africa, Group 3 in Asia and Oceania, Group 6 in Europe, and Group 7 in South America (Table 1). The remaining groups (Groups 1, 4, 5, 8 and 9) did not reach statistical significance, but showed the highest representation in specific regions: Group 1 in Southern Africa, Group 4 in Eastern Africa, Group 5 in Central and West Africa, Group 8 in South and Central America, and Group 9 in the Americas (North America, Central America, Southern America and the West Indies) (Table S5). These results highlight both broad-scale and region-specific geographic patterns among cultivated cowpea, reflecting historical dispersal patterns.

**Table 1.** Non-parametric assessment of geographic enrichment of group proportions for the nine genetic groups. Mean group proportions for each region are presented with compact letter display indicating a significant difference in means across regions. Significant group enrichment for one geographic region are underlined. Note: North and Central America are separated into two regions. *n* = Geographic region sample size.

|  |  | Group 1 |  | Group 2 |  | Group 3 |  | Group 4 |  | Group 5 |  | Group 6 |  | Group 7 |  | Group 8 |  | Group 9 |  |
| --- | --- | --- | --- | --- | --- | --- | --- | --- | --- | --- | --- | --- | --- | --- | --- | --- | --- | --- | --- |
| Region | *n* | Mean | sig | Mean | sig | Mean | sig | Mean | sig | Mean | sig | Mean | sig | Mean | sig | Mean | sig | Mean | sig |
| Africa | 5,125 | 0.152 | a | <u>0.272</u> | <u>a</u> | 0.041 | b | 0.114 | b | 0.154 | a | 0.088 | e | 0.075 | bc | 0.045 | e | 0.059 | c |
| Asia | 2,654 | 0.042 | d | 0.017 | d | <u>0.429</u> | <u>a</u> | 0.117 | b | 0.026 | c | 0.134 | c | 0.115 | b | 0.073 | d | 0.046 | d |
| Central America | 82 | 0.077 | abc | 0.009 | bcd | 0.103 | b | 0.084 | bd | 0.067 | abc | 0.122 | bcde | 0.076 | bc | 0.232 | ab | 0.230 | ab |
| Europe | 255 | 0.019 | f | 0.017 | bc | 0.107 | bc | 0.032 | e | 0.032 | d | <u>0.678</u> | <u>a</u> | 0.065 | d | 0.033 | f | 0.016 | e |
| North America | 968 | 0.039 | e | 0.017 | c | 0.060 | c | 0.066 | c | 0.064 | b | 0.222 | b | 0.096 | c | 0.151 | c | 0.284 | a |
| Oceania | 68 | 0.050 | acd | 0.018 | bd | <u>0.322</u> | <u>a</u> | 0.184 | a | 0.038 | bcd | 0.098 | de | 0.055 | bc | 0.143 | bc | 0.092 | cd |
| South America | 277 | 0.100 | b | 0.020 | bd | 0.044 | b | 0.078 | ad | 0.052 | b | 0.068 | e | <u>0.156</u> | <u>a</u> | 0.280 | a | 0.202 | a |
| Unknown | 180 | 0.053 | cde | 0.025 | bc | 0.133 | b | 0.096 | bc | 0.046 | bc | 0.195 | bcd | 0.102 | bc | 0.178 | bc | 0.173 | b |

#### Phylogeny of cowpea

The cowpea phylogenetic tree (Figure 4) is well supported, with all 9,607 nodes showing bootstrap (BS) values >50% and a mean of 80.7% (Table S6). Support is highest for small terminal groups (BS 90 – 100%), whereas backbone edges are shorter and moderately supported (mean BS 64%). A total of 34 clades were identified along the backbone of the tree (Table S6), 12 of which contain over 200 accessions, including seven clades with more than 500 accessions. A subset of these large clades contain accessions that generally align with the populations, groups and have trends for improvement status, geographic provenance, and cultivar group (Figure S9). A summary of the clades within this tree and detailed historic, taxonomic and geographic patterns can be found in Appendix S1 and Table S6).

**Figure 4.**
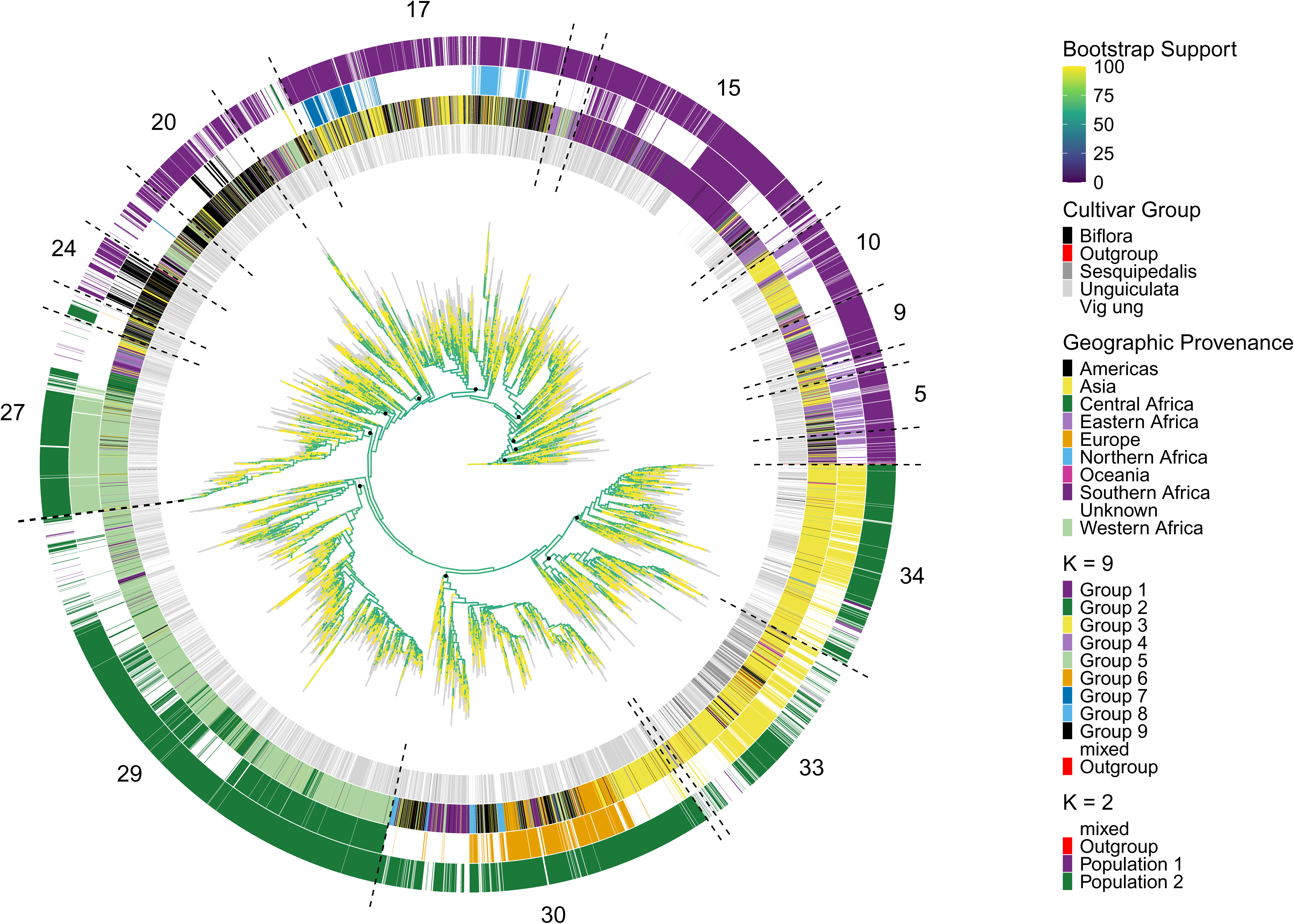
Maximum likelihood phylogeny of 9,610 cowpea accessions implemented in IQ-TREE. The twelve largest clades are numbered outside the tree with dashed lines indicating clade boundaries and black circles indicating their common ancestral node. Bootstrap support values for each node are shown as a continuous color scale on the branches from 0% in dark blue to 100% in yellow, while grey indicates the branches leading to the tips of the tree. Coloured bars on the outside of the tree ordered from the centre ring outwards show: the cultivar group, geographic provenance, genetic group at *K* = 9, and population at *K* = 2 for each accession. See Table S6 for accession names and their corresponding clade.

#### Genetic diversity and relationships between genetic groups

Analysis of molecular variance (AMOVA) was computed between the populations, and groups excluding the mixed accessions. Total molecular variance was partitioned into 32% between populations and 68% within populations at *K* = 2. At *K* = 3, total molecular variance was partitioned into 37% between populations and 63% within populations. At *K* = 9, total molecular variance was partitioned into 50% between groups and 50% within groups (Table 2). Although lower percentages of variation were found between groups at all hierarchy levels, all variance partitioning was statistically significant (*p* < 0.01). The analyses indicated high variance prevailing within the groups identified. AMOVA was also computed between collections and most molecular variation was partitioned within collections (95%) compared to between collections (5%, Table 2).

**Table 2.** Analysis of molecular variance (AMOVA) partitioning the variance between and within genetic groups and collection.

| Hierarchy level | Source of variation | df | SS | MS | p-value | Variance (%) |
| --- | --- | --- | --- | --- | --- | --- |
| Population (*K* = 2) | Between populations | 1 | 975285.8 | 975285.8019 | < 0.001 | 31.95 |
|  | Within populations | 7528 | 4186342.2 | 556.1028 |  | 68.05 |
|  | Total | 7529 | 5161628 | 685.5662 |  | 100 |
| Population (*K* = 3) | Between populations | 2 | 1198696 | 599347.781 | < 0.001 | 36.74 |
|  | Within populations | 6497 | 3338295 | 513.821 |  | 63.26 |
|  | Total | 6499 | 4536991 | 698.106 |  | 100 |
| Group (*K* = 9) | Between groups | 8 | 1205494 | 150686.7596 | < 0.001 | 50.08 |
|  | Within group | 4015 | 1423063 | 354.4365 |  | 49.92 |
|  | Total | 4023 | 2628557 | 653.3822 |  | 100 |
| Collection | Between collections | 6 | 200120.5 | 33353.4246 | < 0.001 | 5.32 |
|  | Within collection | 9602 | 6346552.6 | 660.9615 |  | 94.68 |
|  | Total | 9608 | 6546673.2 | 681.3773 |  | 100 |

df = degrees of freedom; SS = sum of squares; MS = mean square. *p*-value calculated from 999 permutations

Diversity metrics including the number of private alleles, allelic richness, nucleotide diversity, inbreeding co-efficient (F_IS_), IBD, observed heterozygosity and the number of private alleles were assessed for each of the nine groups. Groups 1 (Southern Africa), 2 (Western/Central Africa) and 3 (Asia) contained the most private alleles (*n* = 1998 – 4641, Table S7) compared to Groups 4 (Eastern Africa), 5 (Central/Western Africa), 6 (Europe), 7 (South America), 8 (South/Central America) and 9 (Americas) (*n* = 0 – 656). Group 4 contained the most allelic richness (0.182) and nucleotide diversity (0.16) but a relatively high inbreeding co-efficient (0.93) and few private alleles (*n* = 656, Table 3). Groups 1, 3 and 4 had large π values compared to Groups 7, 6 and 8. Groups 3 – 5 were the most inbred compared to Groups 1, 6 – 8 and Group 5 had the lowest observed heterozygosity (*H*_O_ = 0.007) (Table 3). Group 1 had the lowest inbreeding co-efficient (F_IS_ = 0.77, Table 3), the largest observed heterozygosity (*H*_O_ = 0.033), and a large proportion of breeding material (i.e., *π̂* of ∼ 0.5 is indicative of parent/child relationships and sib-pairs, Figure 5a). Africa was the only continent to contain private alleles (Table S8) and had the largest nucleotide diversity and allelic richness (Table S9).

**Figure 5.**
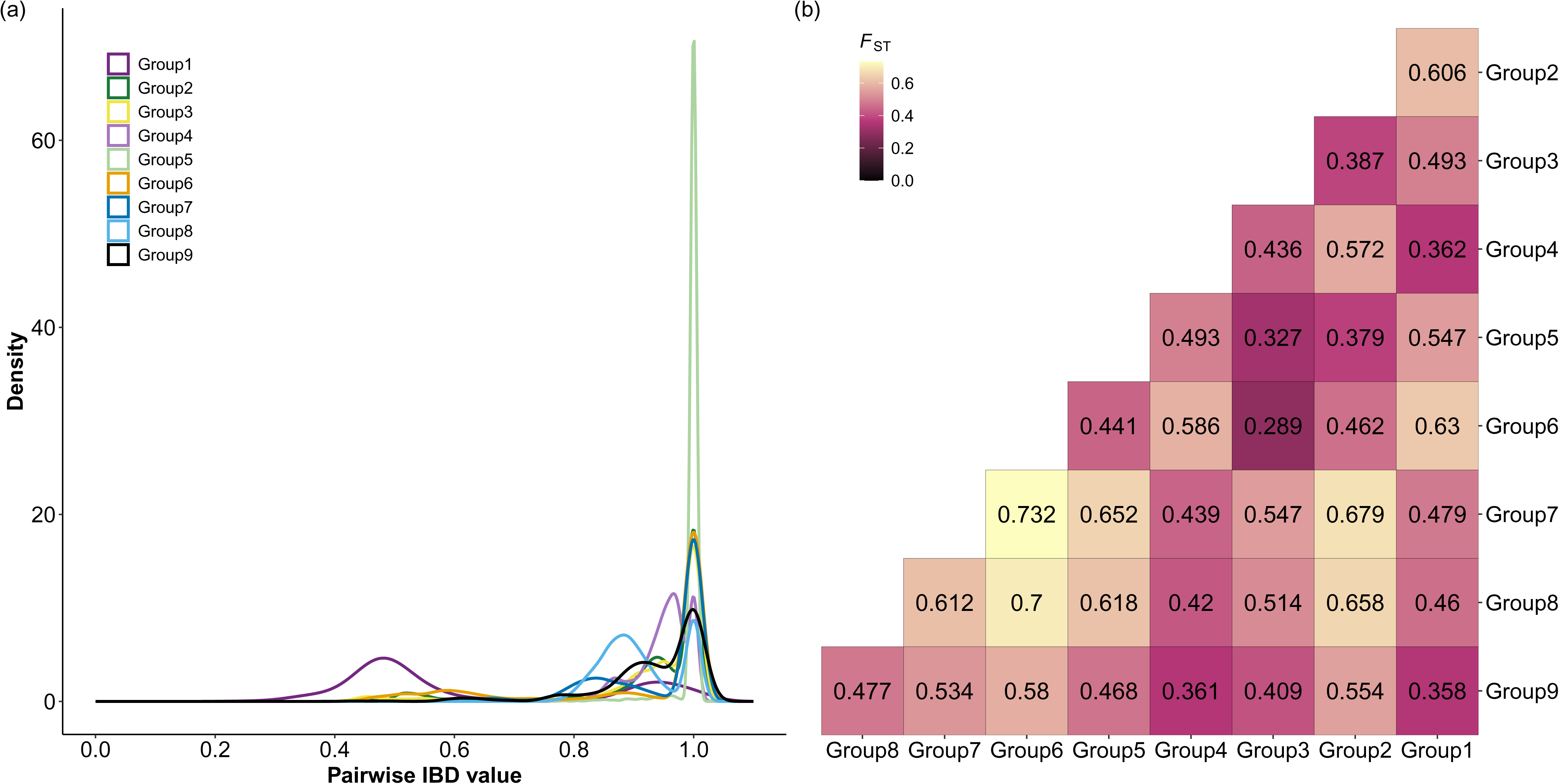
Population relatedness of the non-redundant global cowpea collection. (a) Pairwise identity by descent (IBD) frequency distribution for each genetic group at *K* = 9. (b) Pairwise *F*_ST_ between genetic groups at *K* = 9, coloured continuously from low differentiation (black) to high differentiation (yellow).

**Table 3.** Diversity metrics of the nine genetic groups.

| Group | Allelic richness | Private alleles | π | *F*<sub>IS</sub> | *H*<sub>O</sub> | *H*<sub>E</sub> |
| --- | --- | --- | --- | --- | --- | --- |
| Group 1 (Purple) | 1.72 | 2089 | 0.147 | 0.774 | 0.033 | 0.147 |
| Group 2 (Green) | 1.69 | 4641 | 0.104 | 0.907 | 0.010 | 0.104 |
| Group 3 (Yellow) | 1.74 | 1998 | 0.158 | 0.934 | 0.011 | 0.158 |
| Group 4 (Light Purple) | 1.82 | 656 | 0.161 | 0.932 | 0.011 | 0.160 |
| Group 5 (Light Green) | 1.60 | 630 | 0.108 | 0.937 | 0.007 | 0.108 |
| Group 6 (Orange) | 1.68 | 8 | 0.078 | 0.872 | 0.010 | 0.078 |
| Group 7 (Blue) | 1.73 | 0 | 0.065 | 0.846 | 0.010 | 0.065 |
| Group 8 (Light Blue) | 1.69 | 9 | 0.081 | 0.846 | 0.012 | 0.081 |
| Group 9 (Black) | 1.63 | 0 | 0.134 | 0.924 | 0.010 | 0.134 |

π = Weighted mean nucleotide diversity, *F*<sub>IS</sub> = weighted mean inbreeding co-efficient, *H*<sub>O</sub> = Observed heterozygosity, *H*<sub>E</sub> = Expected heterozygosity

Pairwise *F*_ST_, or the population stratification statistic, was estimated between the *K* = 9 groups (Figure 5b). The highest genetic differentiation of 0.73 was found between Group 6 (Mediterranean) and Group 7 (South American). Similar high differentiation of 0.7 was also observed between Group 6 and Group 8 (Central and South America). High pairwise *F*_ST_ values of 0.679 and 0.658 also indicated that Group 7 and Group 8 were distinct from Group 2 (West African). In contrast, the lowest genetic differentiation of 0.289 was found between Group 6 and Group 3 (Asian).

## DISCUSSION

The purpose of this study was to examine population structure and diversity of cowpea sourced from multiple genebanks and collections. The accessions in this collection reflect the global genetic diversity of extant cowpea. This dataset includes the largest cohort of cowpea genotyped using a single technology to date. The collection includes previously uncharacterised accessions and integrates previously characterised collections, creating a valuable resource for cowpea research and crop improvement.

### Geographic structure of genetic diversity mirrors historical dispersal

Cowpea’s global movement and distribution are mirrored in its genetic structure and diversity. As cowpea was introduced to different regions, selection for local environments and human preference traits shaped its genetic characteristics. The genetic variation observed in cowpea populations today reflect historical migration patterns and regions of cultivation with independent introduction events (See Appendix S1 for detailed information). Previous studies have primarily focused on genetic diversity within African cowpea populations, often including limited samples from Asia and Europe (Herniter et al., 2020, Muñoz-Amatriaín et al., 2021, Fiscus et al., 2024). However, no study to date has comprehensively analysed cowpea representatives from multiple continents to explore their genetic relationships.

West Africa is widely regarded as a centre of diversity for cultivated cowpea, with earlier studies identifying two major gene pools separating Western and Central African cowpea from Eastern and Southern African cowpea (Steele, 1976, Padulosi et al., 1997, Huynh et al., 2013, Fatokun et al., 2018). In this study, we confirmed the presence of two primary gene pools: Population 1, which includes cowpea from West Africa, East Africa, Southern Africa and the Americas; and Population 2, which consists of cowpea primarily from West Africa, Central Africa, Asia and Europe. However, these gene pools likely reflect an underestimation of *K*, with STRUCTURE detecting the largest change in model likelihood (Janes et al., 2017) and may not reflect the dual domestication events previously hypothesised (Huynh et al., 2013). Alternatively, nine gene pools detected by sNMF reflect anthropogenic dispersal and selection/drift and possibly local adaptation. Assessment of the relationship among the wild progenitor of domesticated cowpea, *Vigna unguiculata* subsp. *unguiculata* var. *spontanea* (Schweinf.) Pasquet formerly known as *Vigna unguiculata* ssp. *dekindtiana*, and a representative sampling of African landraces encompassing broad geographic and agroecological diversity, along with phylogenetic analysis using plastome sequences, or haplotype analysis of domestication genes would help elucidate the domestication history of cowpea.

The primary centre of diversity for cowpea remains in sub-Saharan Africa (Table 3), with Group 4 exhibiting the highest genetic diversity compared to other groups and positioned near the root of the phylogenetic tree with short branch lengths (Figure 4), indicating some Group 4 members may represent landraces most closely related to the original domestication gene pool. The phylogenetic analysis revealed a short backbone with moderate support values, indicating limited resolution among the deeper relationships between accessions. This pattern likely reflects the rapid radiation from Africa with many lineages diverging in a short evolutionary timeframe, thus leaving insufficient genetic signal to confidently resolve basal nodes (Whitfield et al., 2007). Low bootstrap values on the backbone nodes also suggest potential complications such as incomplete lineage sorting or historical geneflow, common phenomena in domesticated species, that can obscure clear bifurcating relationships. These factors complicate the reconstruction of deep evolutionary histories and highlight caution when interpreting relationships among major clades. Despite these limitations, the cowpea phylogenetic tree can still provide meaningful insight into more recent divergences, particularly among well-supported smaller clades and terminal branches. These regions can inform patterns of genetic differentiation and can help identify clusters of closely related accessions. To improve the resolution, future studies could incorporate genome-wide genetic data or employ network-based approaches that better accommodate reticulate evolution and admixture. Overall, the observed phylogenetic structure highlights the complexity of the domestication, radiation and breeding history in cowpea and demonstrates clear lineages between cowpea accessions.

The observation that genetic groups correspond closely with geographic origin indicates that spatial factors have significantly influenced the genetic differentiation among accessions. This pattern may reflect limited gene flow between regions, allowing populations to diverge genetically over time due to isolation. It also supports the potential for local adaptation, where environmental pressures such as climate, soil type, or agricultural practices have selected for region-specific traits. A strong geographic basis may also point to historical selection events or drift that occurred independently across regions, leading to distinct genetic lineages (Appendix S1). This geographic structuring is valuable for conservation and breeding, as it highlights reservoirs of unique genetic variation that may be associated with traits beneficial for specific environments or stress conditions. Understanding the geographic distribution of genetic diversity can inform targeted conservation strategies and guide the selection of accessions for breeding programs aimed at improving crop performance under region-specific constraints.

### Modern germplasm sharing and implications for cowpea conservation and management

Global sharing of cowpea germplasm has led to the presence of duplicated accessions within and between collections, highlighting an important challenge in germplasm management. A total of 1,008 duplicate accessions (9.5%) were identified from this global cowpea collection. Compared to other crop species such as barley, cassava and soybean which show 33% (Milner et al., 2019), 26% (Albuquerque et al., 2019) and 23% (Song et al., 2015) duplication respectively, duplication of cowpea accessions is relatively low. However, this study does not encompass all globally available accessions as Genesys currently contains ∼45,500 amalgamated cowpea accessions (Data accessed through Genesys, https://www.genesys-pgr.org/c/cowpea, 2 October 2025). Numerous collections exist worldwide such as the full collection maintained by the International Institute of Tropical Agriculture (IITA, Ibadan, Nigeria, ∼19,000 accessions), the National Bureau of Plant Genetic Resources (NBPGR, New Delhi, India, 3,704 accessions), and the full collection held at the University of California, Riverside (UCR, California, USA, ∼5,000 accessions). If these additional ∼ 28,000 accessions were assessed, we would expect to identify substantially more duplicated entries within and across collections.

Cowpea accessions have been acquired from more than 120 countries worldwide. Re-collecting at the same locations and sharing germplasm among genebanks can result in duplications within and among genebanks. Most duplicate cowpea accessions were identified between collections (74%), indicating material exchange with appropriate documentation. However, a total of 33,252 pairs of accessions were identified as potential duplicates based on molecular data alone. These pairs may either be duplicate accessions but have incomplete or inconsistent passport information, i.e., the same accession with different names have been acquired multiple times, or they could be different accessions but very closely related, i.e., breeding material. Redundancy within collections increases the management workload for curators and may lead to users selecting duplicated accessions if chosen at random. Pairs of highly similar accessions may be attributed to (1) breeders consistently using improved elite lines as parents in crosses in their programs, (2) backcrossing to introduce simply inherited traits, or (3) selection for specific human and agronomic traits limiting the number of parents used in breeding programs. Implementing pedigree metadata for breeding material, updating accession passport descriptors and standardising trait recording will enhance the accessibility of germplasm for breeding programs.

This study revealed that most molecular variance was partitioned within collections (Table 2), rather than between, indicating that local collections contain distinct germplasm. Sequencing technology has reached a point where a global coordinated effort among all genebanks can genetically curate all accessions to find unique accessions within them. This study demonstrates the utility of low-density SNP markers for identifying potentially duplicated accessions. This will enable genebank curators to subset these pairs to investigate duplication status further through either higher resolution genetic sequencing or phenotypic characterisation. Furthermore, new field collections should be genotyped and assessed for similarity to pre-existing accessions to avoid redundancy. These globally unique accessions should then be prioritised and shared with other genebanks for additional backup of those irreplaceable accessions.

Identity by descent analyses provides the first evidence towards identifying duplicate genotypes and pedigree-related relationships. Within accession variation was not assessed in this study, as such variation was expected to be low for cowpea (Xiong et al., 2016, Gumede et al., 2022) due to the low outcrossing rate of 0-5%. Despite this, any within accession variation effects on the estimation of pairwise similarity and duplicate identification cannot be ruled out. Although cowpea has a relatively small genome (640Mbp) (Lonardi et al., 2019), the modest number of genetic markers (3,073 SNPs) used here to identify duplicates may limit duplication detection. Combined, these analyses may not have identified the true level of duplication in the cowpea global collection but do indicate potential candidates for germplasm management. Corroborating evidence of the common origin of two accessions can be identified through (1) comparison of vouchers in the genebanks’ herbarium, (2) inspection of the original (paper) passport records, (3) assessing higher density marker data for a larger number of individuals from candidate duplicates, and (4) field evaluation in which candidate duplicates are grown side by side. Although a daunting task, the time invested in working through the list of duplicates case by case will outweigh the accumulating cost of maintaining superfluous material.

Geographic clustering observed in cowpea populations demonstrates the influence of regional breeding practices and environmental conditions on genetic variation. This highlights the importance of accounting for geographic structure in future analyses, and in future germplasm collection missions. The declining cost of genotyping over the past decade has made advanced genetic tools more accessible, enabling even small-scale projects and genebanks to adopt high-throughput approaches. The identification of duplicate accessions reveals the redundancy that can accumulate over time, posing challenges to the efficient use of germplasm. These findings carry important implications for both scientific research and crop improvement efforts. Duplicate entries contribute to the inefficiencies in germplasm conservation, inflate operational costs, and dilute the uniqueness of genetic resources. Addressing this issue is critical for optimizing cowpea germplasm management and ensuring that valuable genetic diversity is preserved and utilized effectively. Transitioning from paper-based records to digital tracking systems will enhance germplasm management and help reduce duplication across genebanks. Furthermore, updating accession passport descriptors and standardizing trait recording practices will improve the accessibility and utility of germplasm collections for breeding programs, ultimately supporting more targeted and efficient crop improvement strategies.

## ACKNOWLEDGMENTS

We thank Virginia McQueen and Erica Steadman at the Australian Grains Genebank (AGG) for assisting with tissue sampling. We thank Shyam Tallury and the United States Department of Agriculture (USDA) Genebank; Olaniyi Oyatomi and the International Institute of Tropical Agriculture (IITA) Genebank; the National Agriculture and Food Research Organization (NARO) Genebank; Rogerio Chiulele from Eduardo Mondlane University of Mozambique (EMU); the Australian Grains Genebank (AGG); Timothy Close and Maria Munoz-Amatriain for providing UCR germplasm; and Jean-Philippe Vielle-Calzada and his team at Cinvestav Langebio for help collecting and providing cowpea accessions.

## AUTHOR CONTRIBUTIONS

EM, DJ, AC, AMGK designed the project; PM, MMW, AC, MDA, TI, JC, POA, JPVC and SN collected and provided plant materials; TS prepared the sequenced samples; SP, AH and SS performed data analyses; SP wrote the manuscript; EM, DJ, MDA, MMW, SS, AH, AMGK, POA, YT, JC, TI and SN revised the manuscript.

## DATA AVAILABILITY

The data that supports the findings of this study are available at UQ eSpace in the dataset “CowpeaDiversityGeneticData” using the following link: (upon manuscript acceptance), in the supplementary material of this article, and in the following GitHub repository: https://github.com/SofiePearson/CowpeaDiversity.

## COMPETING INTERESTS

The authors declare no competing interests.

## FUNDING

This work was supported in part by the Gates Foundation (Hy-Gain INV-002955). The conclusions and opinions expressed in this work are those of the author(s) alone and shall not be attributed to the Foundation. Under the grant conditions of the Foundation, a Creative Commons Attribution 4.0 License has already been assigned to the Author Accepted Manuscript version that might arise from this submission. Please note works submitted as a preprint have not undergone a peer review process.

## SUPPORTING INFORMATION

**Figure S1.** Heatmap showing the density and distribution of 4,290 SNP markers across the eleven cowpea chromosomes.

**Figure S2.** Principal component (PC) analysis of 10,617 accessions with each panel representing various metadata.

**Figure S3.** Support for the number of ancestral populations (*K*) from the ADMIXTURE analysis where *K* was tested from 1 to 40.

**Figure S4.** Probability support for the number of ancestral populations (*K*) from the STRUCTURE analysis where *K* was tested from 1 to 25. Plots produced from the Evanno method.

**Figure S5.** Cross-entropy support for the number of ancestral populations (*K*) from the sNMF analysis where *K* was tested from 1 to 25.

**Figure S6.** Geographic distribution of the Populations at *K* = 2.

**Figure S7.** Geographic distribution of the Populations at *K* = 3.

**Figure S8.** Principal component (PC) analysis of the 9,609 cowpea accessions coloured by group (*K* = 9).

**Figure S9.** Maximum likelihood phylogeny of 9,610 cowpea accessions implemented in IQ-TREE.

**Figure S10.** Maximum likelihood phylogeny of 894 cowpea accessions from Clade 15.

**Figure S11.** Maximum likelihood phylogeny of 1,056 cowpea accessions from Clade 17.

**Figure S12.** Maximum likelihood phylogeny of 384 cowpea accessions from Clade 20.

**Figure S13.** Maximum likelihood phylogeny of 733 cowpea accessions from Clade 27.

**Figure S14.** Maximum likelihood phylogeny of 1,874 cowpea accessions from Clade 29.

**Figure S15.** Maximum likelihood phylogeny of 1,241 cowpea accessions from Clade 30.

**Figure S16.** Maximum likelihood phylogeny of 712 cowpea accessions from Clade 33.

**Figure S17.** Maximum likelihood phylogeny of 748 cowpea accessions from Clade 34.

**Table S1.** Passport descriptors of 10,618 cowpea accessions.

**Table S2.** Percentage of Improvement Status category assignment for 10,617 cultivated cowpea accessions for each collection source.

**Table S3.** Paired potential duplicate accessions (33,252) and paired duplicate accessions (1,329) identified from genetic distance (GD), genomic relatedness (GRM), whole genome identity by descent (PI HAT, Z0, Z1 and Z2), and accession name and alias matching.

**Table S4.** Admixture proportion assignment of each accession to three values of *K*.

**Table S5.** Non-parametric assessment of regional geographic enrichment of group proportions for nine genetic groups.

**Table S6.** Tree data for the circular 9,610 cowpea tree including node, branch lengths, bootstrap support and accession order.

**Table S7.** Counts of private alleles within each genetic group at *K* = 9.

**Table S8.** Counts of private alleles within each continent.

**Table S9.** Diversity metrics for each continent.

**Table S10.** Predicted group membership of 1,008 redundant cowpea accessions at three levels of *K*.

**Table S11.** Number of accessions within each clade by their population assignment at *K* = 2.

**Table S12.** Number of accessions within each clade by their population assignment at *K* = 3.

**Table S13.** Number of accessions within each clade by their group assignment at *K* = 9.

**Table S14.** Percentage of accessions within each clade by their geographic provenance.

**Table S15.** Percentage of accessions within each clade by their improvement status.

**Table S16.** Percentage of accessions within each clade by their collection source.

**Table S17.** Percentage of accessions within each clade by their cultivar group.

**Table S18.** Regional geographic locations for American (Northern, Central and Southern) accessions and their corresponding clade.

**Appendix S1.** Summary of the cowpea phylogenetic tree and geographic relationships.

## Notes

### Competing Interest Statement

The authors have declared no competing interest.

