## Supplemental Figure Captions for "Scaling up orphan crop research: A global genetic perspective of cowpea (*Vigna unguiculata*) diversity from 10,617 accessions"

**Supplementary Figure Captions**

**Figure S1.** Heatmap showing the density and distribution of 4,290 SNP markers across the eleven cowpea chromosomes.

**Figure S2.** Principal component (PC) analysis of 10,617 accessions with each panel representing various metadata.

(a) Cultivar group: black triangles = biflora, grey squares = sesquipedalis, grey plus = unguiculata, and grey circles = species level *Vigna unguiculata* or subspecies *unguiculata*.

(b) – (f) Improvement Status: blue circles = Advanced/Improved Cultivar, Breeding Line, Landrace/Traditional Cultivar, Wild, Weedy. Grey circles = all other lines.

(g) – (m) Collection: orange circles = Australian Grains Genebank (AGG), Instituto de Investigação Agrária de Moçambique (IIAM), International Institute of Tropical Agriculture (IITA), Langebio (LAN), National Agriculture and Food Research Organization (NARO), University of California, Riverside (UCR), United States Department of Agriculture (USDA). Grey circles = all other lines.

(n) – (x) Geographic Provenance: pink circles = Northern Africa, Western Africa, Central Africa, Eastern Africa, Southern Africa, Asia, Northern America, Central America, Southern America, Europe, Oceania. Grey circles = all other lines.

**Figure S3.** Support for the number of ancestral populations (*K*) from the ADMIXTURE analysis where *K* was tested from 1 to 40.

(a) Cross-validation values and standard error from 10 independent runs for *K* = 1 to *K* = 40.

(b) Loglikelihood values and standard error for *K* = 1 to *K* = 40.

(c) Delta *K* for *K* = 2 to *K* = 40. The *K* = 2 value with the greatest support is highlighted in blue.

**Figure S4.** Probability support for the number of ancestral populations (*K*) from the STRUCTURE analysis where *K* was tested from 1 to 25. Plots produced from the Evanno method.

**Figure S5.** Cross-entropy support for the number of ancestral populations (*K*) from the sNMF analysis where *K* was tested from 1 to 25.

(a) Cross-entropy boxplots from 20 iterations for *K* values from 1 – 25.

(b) Change in cross-entropy values with *K* = 2, *K* = 3 and *K* = 9 highlighted in blue.

**Figure S6.** Geographic distribution of the Populations at *K* = 2. For each country, the pie represents the proportion of the ancestral allele frequencies. Purple = Population 1 and Green = Population 2.

**Figure S7.** Geographic distribution of the Populations at *K* = 3. For each country, the pie represents the proportion of the ancestral allele frequencies. Purple = Population 1, Green = Population 2 and Yellow = Population 3.

**Figure S8.** Principal component (PC) analysis of the 9,609 cowpea accessions coloured by group (*K* = 9).

(a) PC1 vs PC2.

(b) PC1 vs PC3.

(c) PC2 vs PC3.

**Figure S9.** Maximum likelihood phylogeny of 9,610 cowpea accessions implemented in IQ-TREE. The 34 clades are highlighted with black circles indicating their common ancestral node. The twelve largest clades are labelled to the right of the tree. Bootstrap support values for each node are shown as a continuous colour scale on the branches from 0% in dark blue to 100% in yellow, while grey indicates the branches leading to the tips of the tree. Taxa names are not presented. Coloured bars on the right-hand side of the tree, ordered from the left to the right show: the improvement status, cultivar group, geographic provenance, genetic group at *K* = 9, population at *K* = 3, population at *K* = 2 and collection source for each accession. Colour keys are presented to the right of the bars. See Table S6 for accession names and their corresponding clade.

**Figure S10.** Maximum likelihood phylogeny of 894 cowpea accessions from Clade 15. The six subclades are highlighted with black circles indicating their common ancestral node. Clades consisting of accessions from common countries are labelled to the right of the tree. Bootstrap support values for each node are shown as a continuous colour scale on the branches from 0% in dark blue to 100% in yellow, while grey indicates the branches leading to the tips of the tree. Three letter country code of the country of provenance is presented for each accession at the tip. Coloured bars on the right-hand side of the tree, ordered from the left to the right show: the improvement status, cultivar group, geographic provenance, and genetic group at *K* = 9 for each accession. Colour keys are presented to the right of the bars. See Table S6 for accession names and their corresponding subclade and order in the tree.

**Figure S11.** Maximum likelihood phylogeny of 1,056 cowpea accessions from Clade 17. The two subclades are highlighted with black circles indicating their common ancestral node. Clades consisting of accessions from common countries or geographic regions are labelled to the right of the tree. Bootstrap support values for each node are shown as a continuous colour scale on the branches from 0% in dark blue to 100% in yellow, while grey indicates the branches leading to the tips of the tree. Three letter country code of the country of provenance is presented for each accession at the tip. Coloured bars on the right-hand side of the tree, ordered from the left to the right show: the improvement status, cultivar group, geographic provenance, and genetic group at *K* = 9 for each accession. Colour keys are presented to the right of the bars. See Table S6 for accession names and their corresponding subclade and order in the tree.

**Figure S12.** Maximum likelihood phylogeny of 384 cowpea accessions from Clade 20. The three subclades are highlighted with black circles indicating their common ancestral node. Clades consisting of accessions from common countries or geographic regions are labelled to the right of the tree. Bootstrap support values for each node are shown as a continuous colour scale on the branches from 0% in dark blue to 100% in yellow, while grey indicates the branches leading to the tips of the tree. Three letter country code of the country of provenance is presented for each accession at the tip. Coloured bars on the right-hand side of the tree, ordered from the left to the right show: the improvement status, cultivar group, geographic provenance, and genetic group at *K* = 9 for each accession. Colour keys are presented to the right of the bars. See Table S6 for accession names and their corresponding subclade and order in the tree.

**Figure S13.** Maximum likelihood phylogeny of 733 cowpea accessions from Clade 27. The four subclades are highlighted with black circles indicating their common ancestral node. Clades consisting of accessions from common countries or geographic regions are labelled to the right of the tree. Bootstrap support values for each node are shown as a continuous colour scale on the branches from 0% in dark blue to 100% in yellow, while grey indicates the branches leading to the tips of the tree. Three letter country code of the country of provenance is presented for each accession at the tip. Coloured bars on the right-hand side of the tree, ordered from the left to the right show: the improvement status, cultivar group, geographic provenance, and genetic group at *K* = 9 for each accession. Colour keys are presented to the right of the bars. See Table S6 for accession names and their corresponding subclade and order in the tree.

**Figure S14.** Maximum likelihood phylogeny of 1,874 cowpea accessions from Clade 29. The three subclades are highlighted with black circles indicating their common ancestral node. Clades consisting of accessions from common countries are labelled to the right of the tree. Bootstrap support values for each node are shown as a continuous colour scale on the branches from 0% in dark blue to 100% in yellow, while grey indicates the branches leading to the tips of the tree. Three letter country code of the country of provenance is presented for each accession at the tip. Coloured bars on the right-hand side of the tree, ordered from the left to the right show: the improvement status, cultivar group, geographic provenance, and genetic group at *K* = 9 for each accession. Colour keys are presented to the right of the bars. See Table S6 for accession names and their corresponding subclade and order in the tree.

**Figure S15.** Maximum likelihood phylogeny of 1,241 cowpea accessions from Clade 30. The 11 subclades are highlighted with black circles indicating their common ancestral node. Clades consisting of accessions from common countries or geographic regions are labelled to the right of the tree. Bootstrap support values for each node are shown as a continuous colour scale on the branches from 0% in dark blue to 100% in yellow, while grey indicates the branches leading to the tips of the tree. Three letter country code of the country of provenance is presented for each accession at the tip. Coloured bars on the right-hand side of the tree, ordered from the left to the right show: the improvement status, cultivar group, geographic provenance, and genetic group at *K* = 9 for each accession. Colour keys are presented to the right of the bars. See Table S6 for accession names and their corresponding subclade and order in the tree.

**Figure S16.** Maximum likelihood phylogeny of 712 cowpea accessions from Clade 33. The two subclades are highlighted with black circles indicating their common ancestral node. Clades consisting of accessions from common countries are labelled to the right of the tree and yardlong (sesquipedalis) accessions are highlighted in grey boxes. Bootstrap support values for each node are shown as a continuous colour scale on the branches from 0% in dark blue to 100% in yellow, while grey indicates the branches leading to the tips of the tree. Three letter country code of the country of provenance is presented for each accession at the tip. Coloured bars on the right-hand side of the tree, ordered from the left to the right show: the improvement status, cultivar group, geographic provenance, and genetic group at *K* = 9 for each accession. Colour keys are presented to the right of the bars. See Table S6 for accession names and their corresponding subclade and order in the tree.

**Figure S17.** Maximum likelihood phylogeny of 748 cowpea accessions from Clade 34. The seven subclades are highlighted with black circles indicating their common ancestral node. Clades consisting of accessions from common countries are labelled to the right of the tree. Bootstrap support values for each node are shown as a continuous colour scale on the branches from 0% in dark blue to 100% in yellow, while grey indicates the branches leading to the tips of the tree. Three letter country code of the country of provenance is presented for each accession at the tip. Coloured bars on the right-hand side of the tree, ordered from the left to the right show: the improvement status, cultivar group, geographic provenance, and genetic group at *K* = 9 for each accession. Colour keys are presented to the right of the bars. See Table S6 for accession names and their corresponding subclade and order in the tree.
