## Supplemental Figure 1 for "Scaling up orphan crop research: A global genetic perspective of cowpea (*Vigna unguiculata*) diversity from 10,617 accessions"

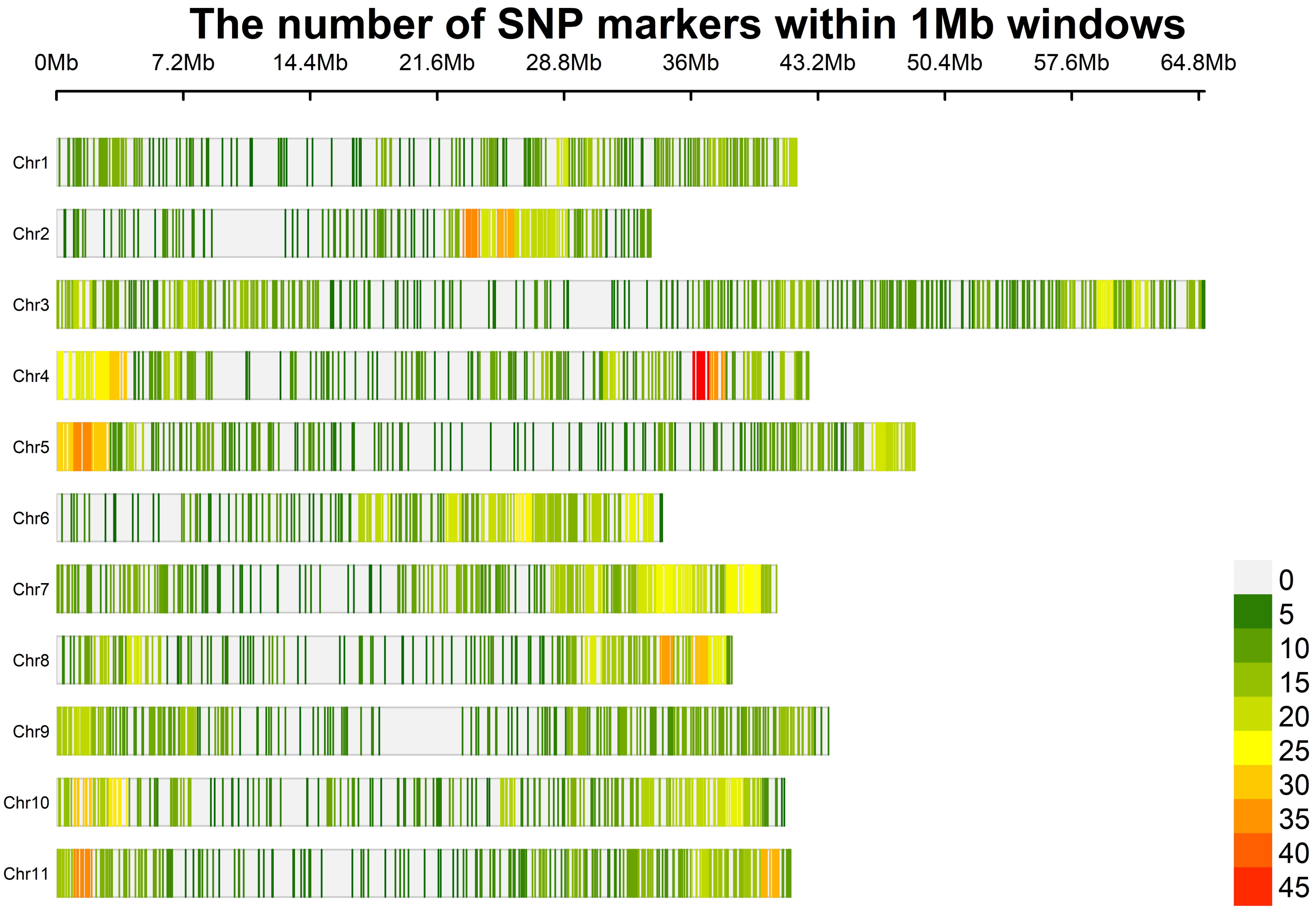


**Figure S1.** Heatmap showing the density and distribution of 4,290 SNP markers across the eleven cowpea chromosomes.
