## Supplemental Figure 2 for "Scaling up orphan crop research: A global genetic perspective of cowpea (*Vigna unguiculata*) diversity from 10,617 accessions"

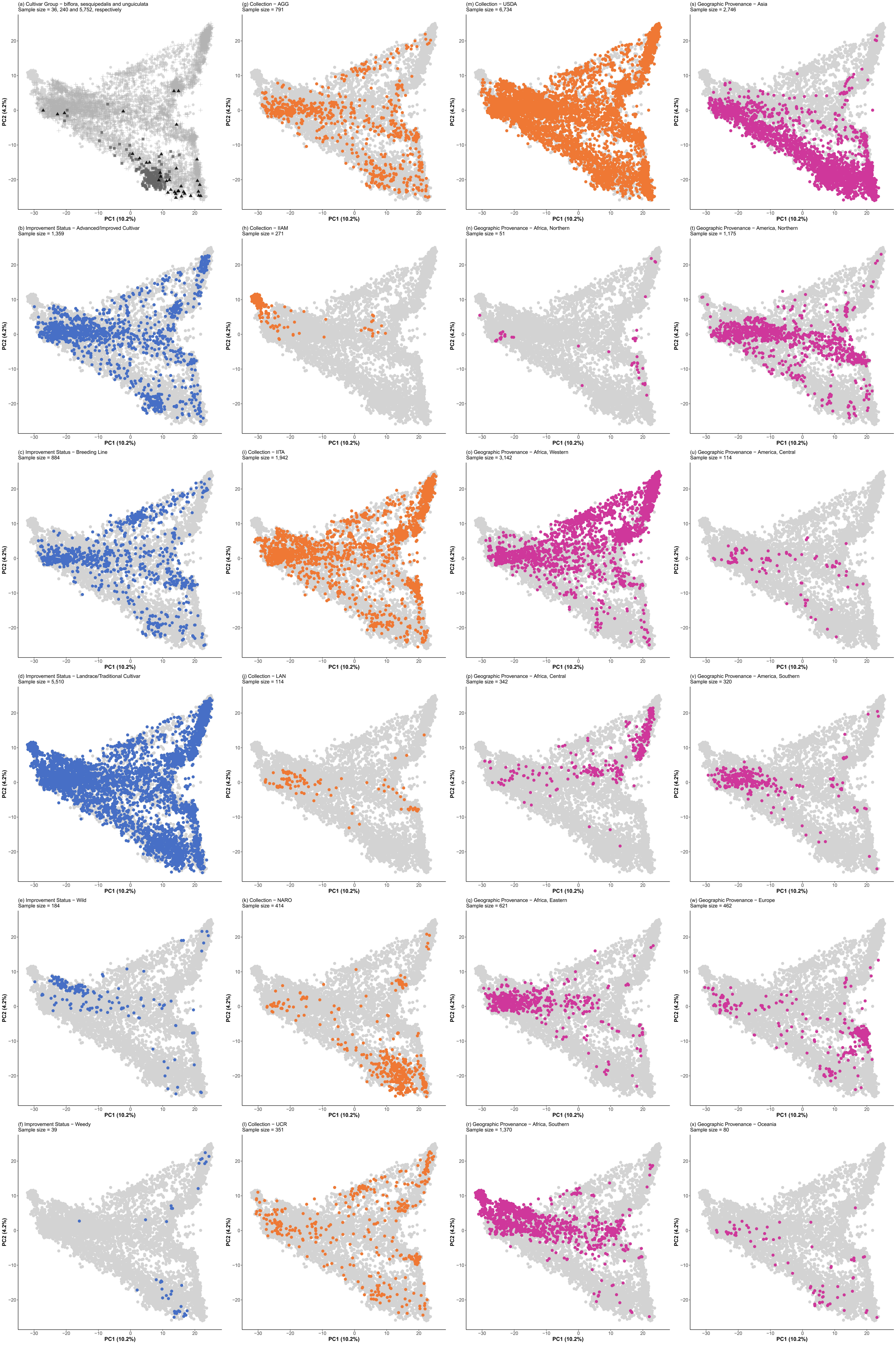


**Figure S2.** Principal component (PC) analysis of 10,617 accessions with each panel representing various metadata.
(a) Cultivar group: black triangles = biflora, grey squares = sesquipedalis, grey plus = unguiculata, and grey circles = species level *Vigna unguiculata* or subspecies *unguiculata*.
(b) – (f) Improvement Status: blue circles = Advanced/Improved Cultivar, Breeding Line, Landrace/Traditional Cultivar, Wild, Weedy. Grey circles = all other lines.
(g) – (m) Collection: orange circles = Australian Grains Genebank (AGG), Instituto de Investigação Agrária de Moçambique (IIAM), International Institute of Tropical Agriculture (IITA), Langebio (LAN), National Agriculture and Food Research Organization (NARO), University of California, Riverside (UCR), United States Department of Agriculture (USDA). Grey circles = all other lines.
(n) – (x) Geographic Provenance: pink circles = Northern Africa, Western Africa, Central Africa, Eastern Africa, Southern Africa, Asia, Northern America, Central America, Southern America, Europe, Oceania. Grey circles = all other lines.
