## Supplemental Figure 3 for "Scaling up orphan crop research: A global genetic perspective of cowpea (*Vigna unguiculata*) diversity from 10,617 accessions"

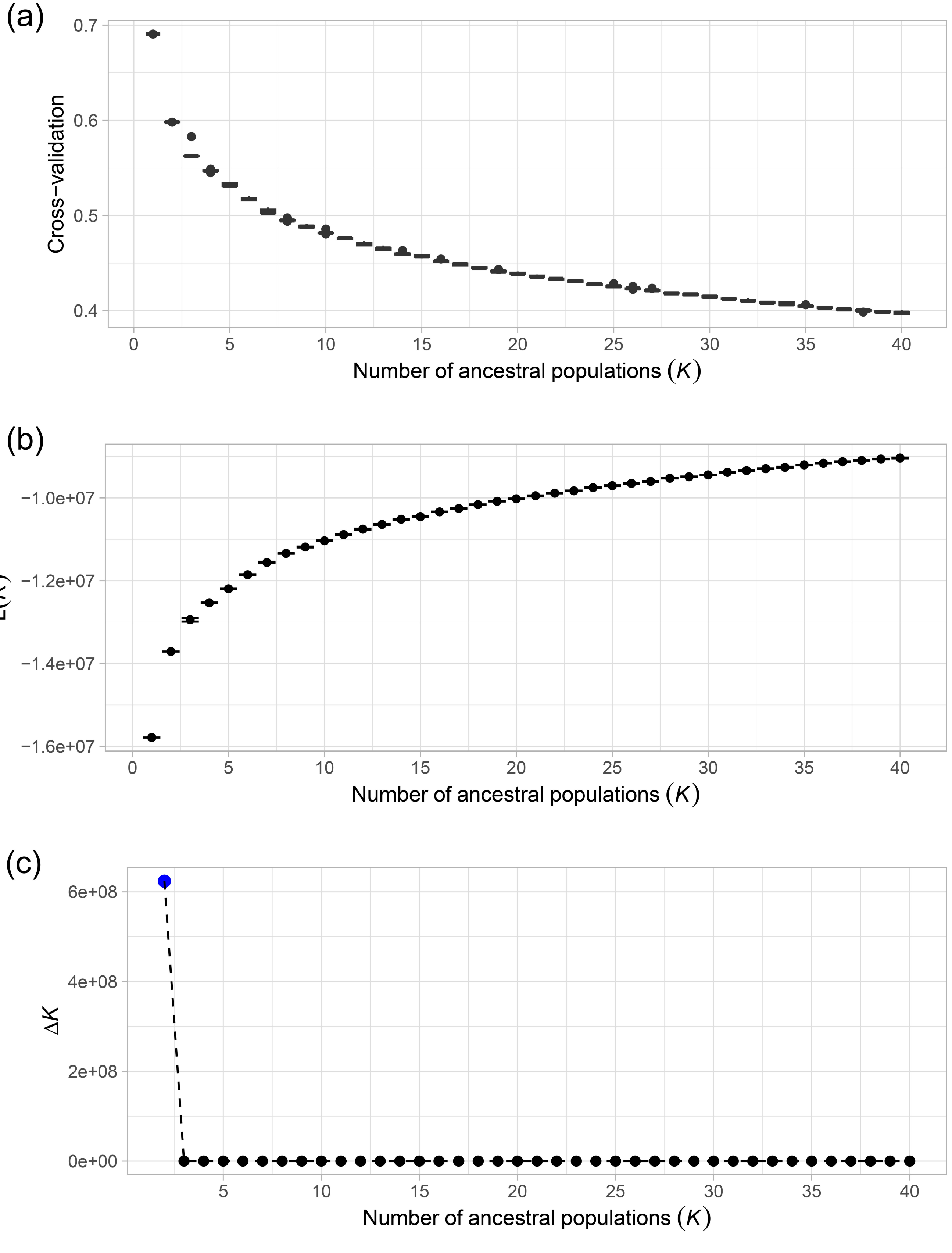


**Figure S3.** Support for the number of ancestral populations (*K*) from the ADMIXTURE analysis where *K* was tested from 1 to 40.
(a) Cross-validation values and standard error from 10 independent runs for *K* = 1 to *K* = 40.
(b) Loglikelihood values and standard error for *K* = 1 to *K* = 40.
(c) Delta *K* for *K* = 2 to *K* = 40. The *K* = 2 value with the greatest support is highlighted in blue.
