## Supplemental Figure 4 for "Scaling up orphan crop research: A global genetic perspective of cowpea (*Vigna unguiculata*) diversity from 10,617 accessions"

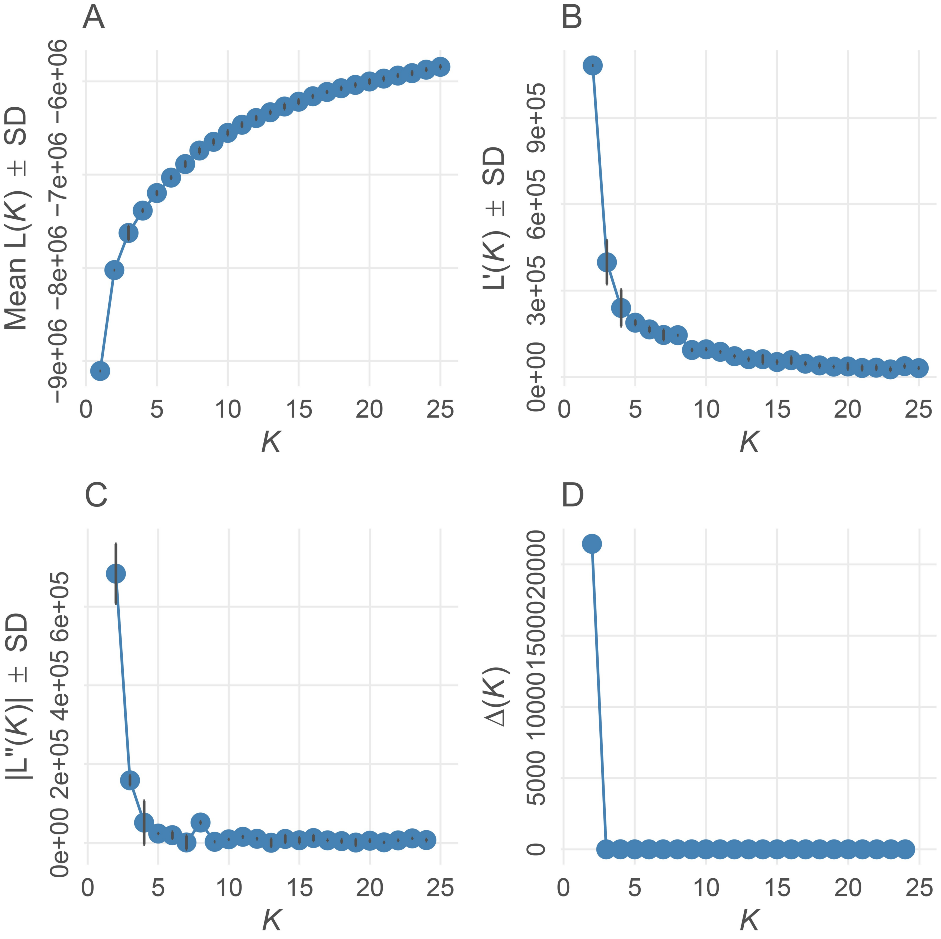


**Figure S4.** Probability support for the number of ancestral populations (*K*) from the STRUCTURE analysis where *K* was tested from 1 to 25. Plots produced from the Evanno method.
