## Supplemental Figure 5 for "Scaling up orphan crop research: A global genetic perspective of cowpea (*Vigna unguiculata*) diversity from 10,617 accessions"

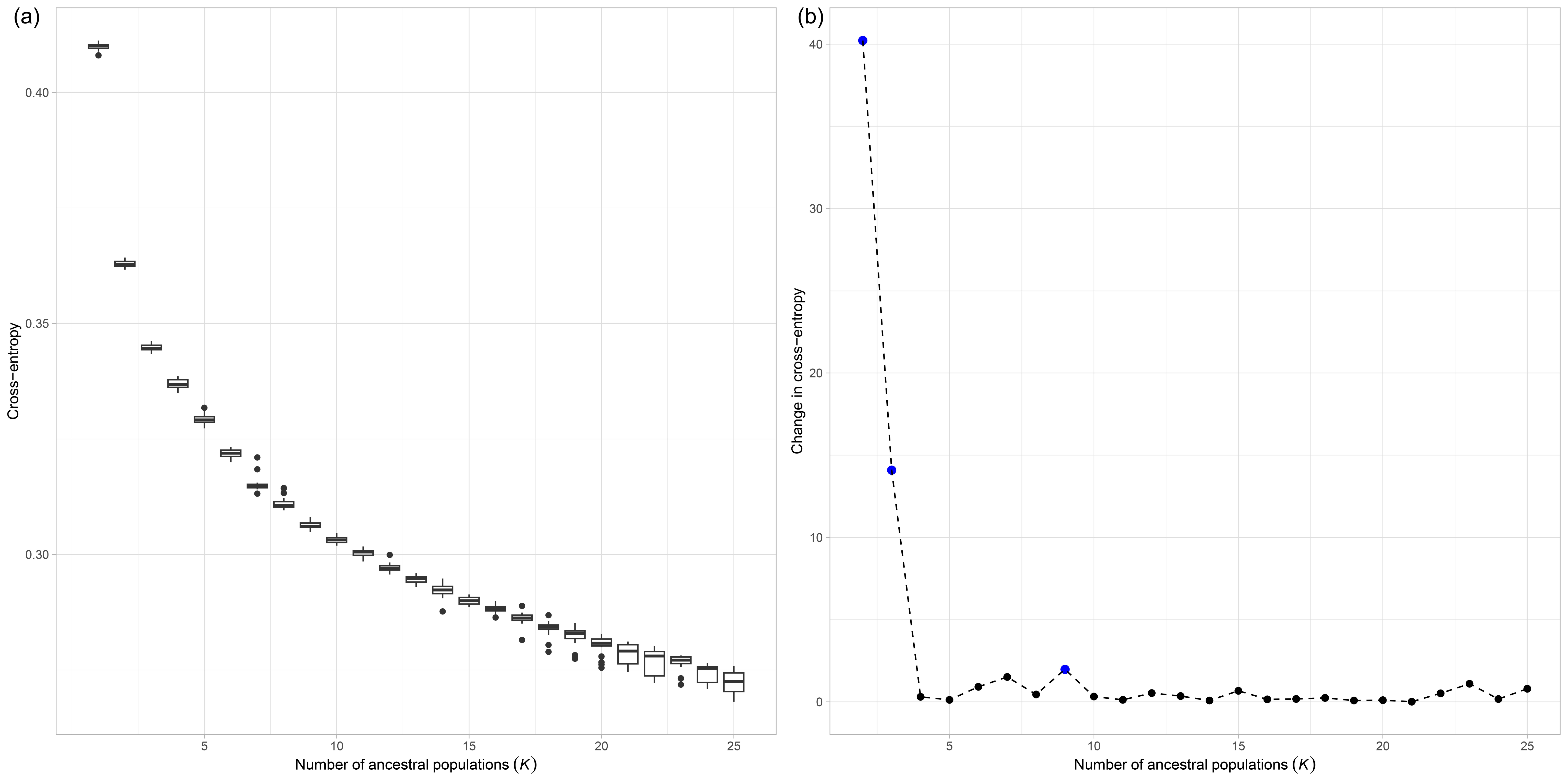


**Figure S5.** Cross-entropy support for the number of ancestral populations (*K*) from the sNMF analysis where *K* was tested from 1 to 25.
(a) Cross-entropy boxplots from 20 iterations for *K* values from 1 – 25.
(b) Change in cross-entropy values with $K = 2$, $K = 3$ and $K = 9$ highlighted in blue.
