## Supplemental Figure 6 for "Scaling up orphan crop research: A global genetic perspective of cowpea (*Vigna unguiculata*) diversity from 10,617 accessions"

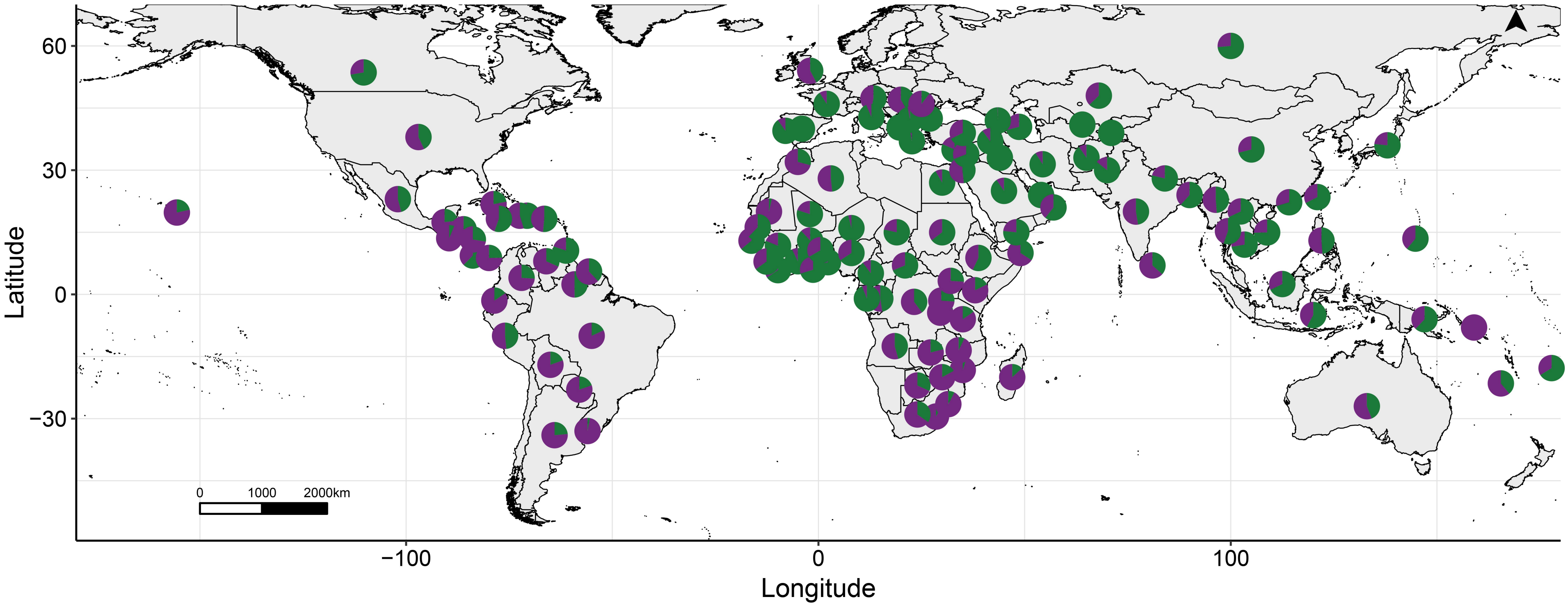


**Figure S6.** Geographic distribution of the Populations at *K* = 2. For each country, the pie represents the proportion of the ancestral allele frequencies. Purple = Population 1 and Green = Population 2.
