## Supplemental Figure 7 for "Scaling up orphan crop research: A global genetic perspective of cowpea (*Vigna unguiculata*) diversity from 10,617 accessions"

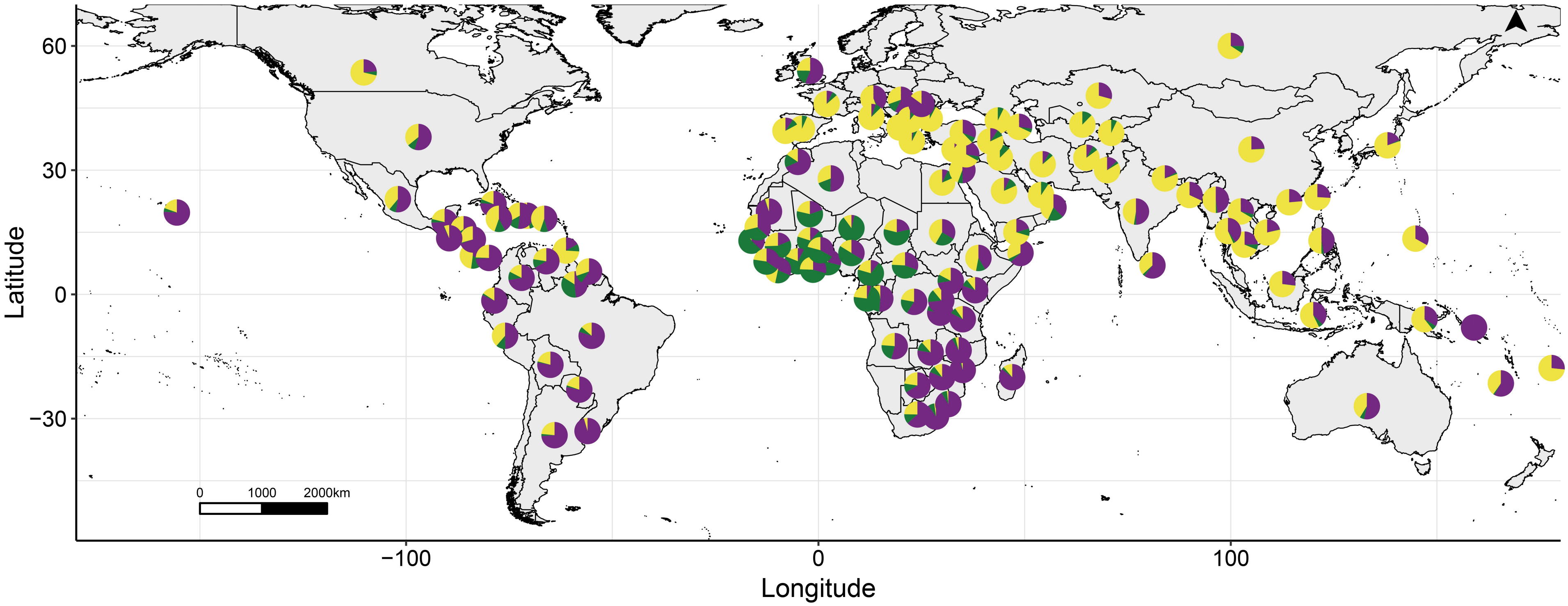


**Figure S7.** Geographic distribution of the Populations at *K* = 3. For each country, the pie represents the proportion of the ancestral allele frequencies. Purple = Population 1, Green = Population 2 and Yellow = Population 3.
