## Supplemental Figure 8 for "Scaling up orphan crop research: A global genetic perspective of cowpea (*Vigna unguiculata*) diversity from 10,617 accessions"

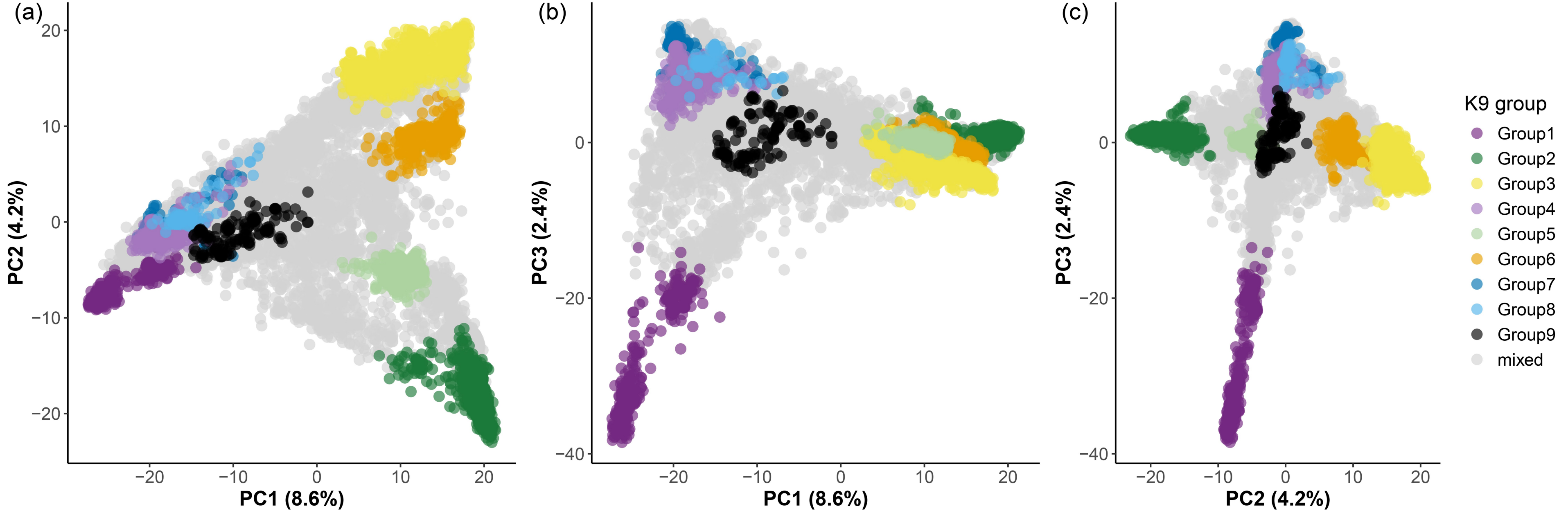


**Figure S8.** Principal component (PC) analysis of the 9,609 cowpea accessions coloured by group (*K* = 9).
(a) PC1 vs PC2.
(b) PC1 vs PC3.
(c) PC2 vs PC3.
