## Supplemental Figure 9 for "Scaling up orphan crop research: A global genetic perspective of cowpea (*Vigna unguiculata*) diversity from 10,617 accessions"

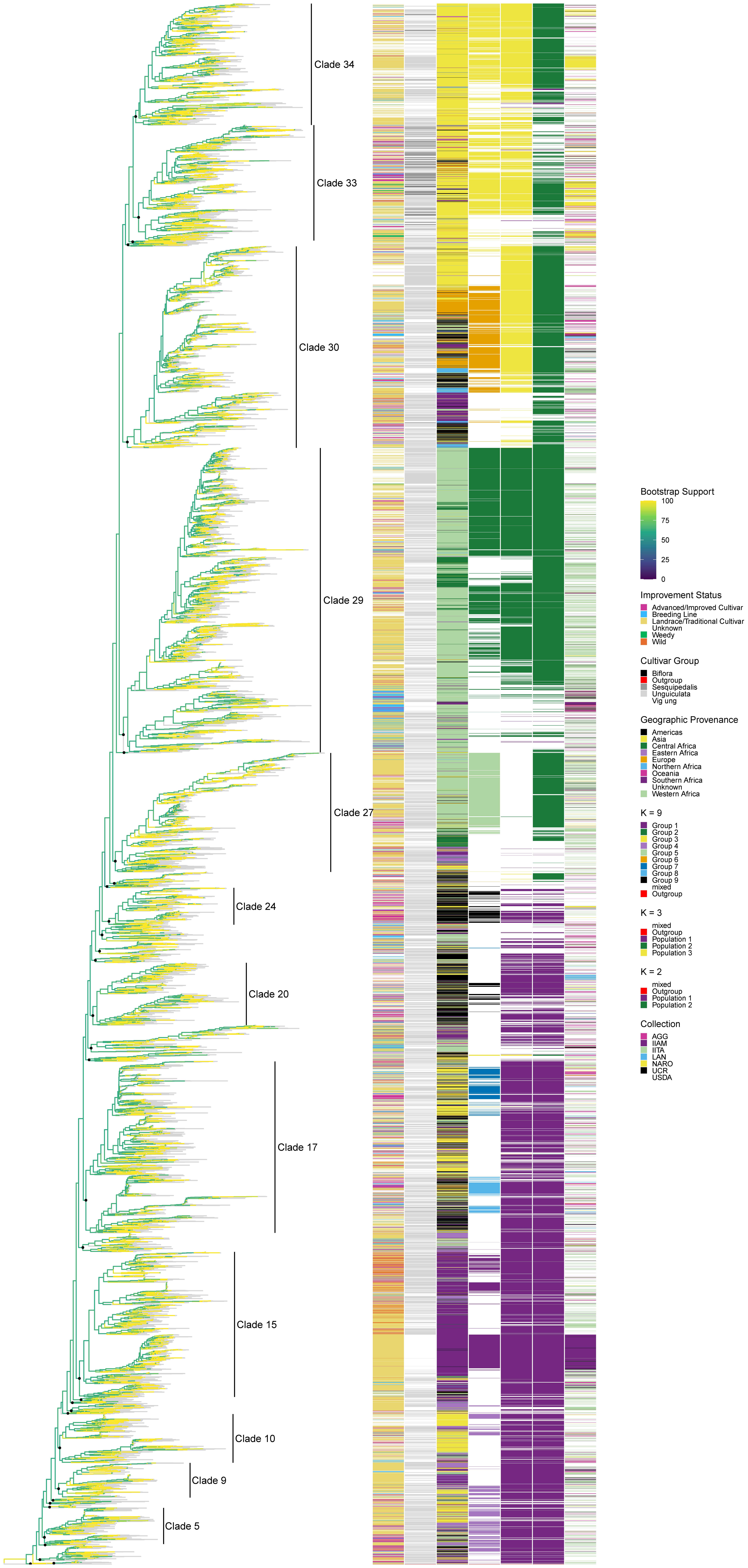


**Figure S9.** Maximum likelihood phylogeny of 9,610 cowpea accessions implemented in IQ-TREE. The 34 clades are highlighted with black circles indicating their common ancestral node. The twelve largest clades are labelled to the right of the tree. Bootstrap support values for each node are shown as a continuous colour scale on the branches from 0% in dark blue to 100% in yellow, while grey indicates the branches leading to the tips of the tree. Taxa names are not presented. Coloured bars on the right-hand side of the tree, ordered from the left to the right show: the improvement status, cultivar group, geographic provenance, genetic group at $K = 9$, population at $K = 3$, population at $K = 2$ and collection source for each accession. Colour keys are presented to the right of the bars. See **Table S6** for accession names and their corresponding clade.
