## Supplemental Figure 17 for "Scaling up orphan crop research: A global genetic perspective of cowpea (*Vigna unguiculata*) diversity from 10,617 accessions"

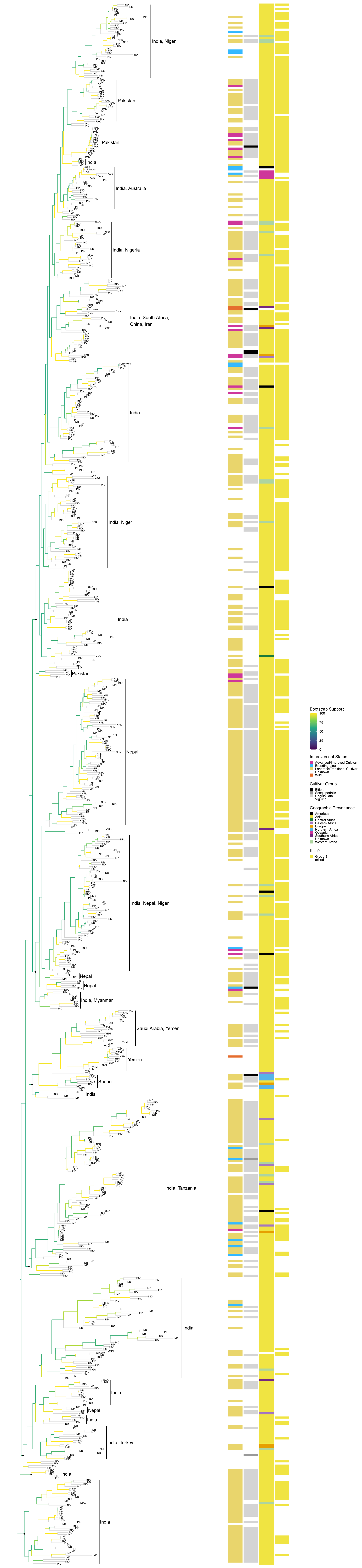


**Figure S17.** Maximum likelihood phylogeny of 748 cowpea accessions from Clade 34. The seven subclades are highlighted with black circles indicating their common ancestral node. Clades consisting of accessions from common countries are labelled to the right of the tree. Bootstrap support values for each node are shown as a continuous colour scale on the branches from 0% in dark blue to 100% in yellow, while grey indicates the branches leading to the tips of the tree. Three letter country code of the country of provenance is presented for each accession at the tip. Coloured bars on the right-hand side of the tree, ordered from the left to the right show: the improvement status, cultivar group, geographic provenance, and genetic group at $K = 9$ for each accession. Colour keys are presented to the right of the bars. See **Table S6** for accession names and their corresponding subclade and order in the tree.
