## Supplemental Appendix 1 for "Scaling up orphan crop research: A global genetic perspective of cowpea (*Vigna unguiculata*) diversity from 10,617 accessions"

**Appendix S1. Supplementary material**

From: Sofie M. Pearson, Adrian Hathorn, Shichao Sun, Alan Cruickshank, Tracey L. Shatte, Paulino Munisse, Mercy Macharia Wairimu, Joann Conner, Anna M.G. Koltunow, Jean-Philippe Vielle-Calzada, Peggy Ozias-Akins, Takayoshi Ishii, Matteo Dell Acqua, Sally Norton, Yongfu Tao, David Jordan and Emma Mace. Scaling up orphan crop research: A global genetic perspective of cowpea (*Vigna unguiculata*) diversity from 10,617 accessions.

**Summary of the cowpea phylogenetic tree and geographic relationships**

Overall, the cowpea phylogenetic tree (Figure 4) is well supported, with all nodes showing bootstrap (BS) values >50% and a mean of 80.7% (Table S6). Support is highest for small terminal groups (BS 90-100%), whereas backbone branches are shorter and moderately supported (mean BD 64%). The tree can be roughly divided into twelve larger clades which generally align with the genetic populations at *K* = 2, 3 and 9,, and have trends for geographic provenance, improvement status, collection, and cultivar group (Figure S9). Trends across the clades are presented below. Additionally, geographic, historic and taxonomic patterns are presented to build upon known geographic spread of cowpea.

<u>Populations (*K* = 2)</u>

At *K* = 2, Clades 1 – 16, 20 – 22, and 25 collectively contain 2,230 accessions from Population 1 (65%); Clades 26, 30 and 32 collectively contain 1,131 accessions from Population 2 (27.5%); and clades 17 – 19, 23, 24, 27, 29, 31, 33 and 34 contain accessions from both Populations 1 and 2. However, Clades 17, 19, 23 and 24 contain more Population 1 accessions (92.3 – 99.8%) than Population 2 accessions (0.2 – 7.7%), and Clades 27, 29, 33 and 34 contain more Population 2 accessions (97.2 – 99.5%) than Population 1 accessions (0.5 – 6.4%, Table S11).

<u>Populations (*K* = 3)</u>

At *K* = 3, Clades 2 – 16, and 19 – 25 collectively contain 2,355 accessions from Population 1 (71.9%); Clades 26, 30 – 32 collectively contain 875 accessions from Population 3 (44.7%); Clades 17, 18, 33 and 34 contain accessions from both Populations 1 and 3; Clade 29 contains accessions from both Populations 1 and 2; and Clade 27 contains few accessions from all three Populations (*n* = 1 – 5). However, Clade 17 contains more Population 1 accessions (*n* = 891) than Population 3 accessions (*n* = 3); Clade 29 contains more Population 2 accessions (*n* = 1264) compared to Population 1 (*n* = 7); and Clades 33 and 34 contain more Population 3 accessions (*n* = 460 and 613, respectively) than Population 1 accessions (*n* = 5 and 6, respectively; Table S12).

<u>Groups (*K* = 9)</u>

At *K* = 9, Clades 3 – 12 and 16 contain Group 4 accessions (*n* = 2 – 150, 0.6 – 42.5%); Clade 15 contains all Group 1 accessions (*n* = 350); Clade 17 contains all Group 8 accessions (*n* = 127) and 98% of Group 7 accessions (*n* = 167); Clade 18 contains six Group 3 accessions (0.7%); Clades 20 and 24 contain 33.3% (*n* = 49) and 66% (*n* = 97), respectively, of Group 9 accessions; Clade 27 contains all Group 5 accessions (*n* = 465); Clade 29 contains all Group 2 accessions (*n* = 976); Clade 30 contains 99.8% of Group 6 accessions (*n* = 531) and 1.4% of Group 3 accessions (*n* = 13); and Clades 33 and 34 contain 47.2% (*n* = 427) and 50.4% (*n* = 456), respectively, of Group 3 accessions (Table S13).

### Geographic provenance

The outgroup accession is from Southern Africa and the sister accession is from West Africa. Clade 1 contains three Indian accessions. Clade 2 contains East African accessions (58%). Clade 3 contains American (31%), East African (27%), West African (15%) and Asian accessions (15%). Clade 4 contains only East African accessions (100%). Clade 5 contains Asian (36%), East African (27%) and West African accessions (16%). Clade 6 contains Asian (41%), Oceanian (14%) and American accessions (18%). Clade 7 contains only Asian (68%) and West African accessions (16%). Clade 8 contains Asian (36%), West African (27%) and American accessions (18%). Clade 9 contains East African (36%), Southern African (29%), Asian (20%) and West African accessions (11%). Clade 10 contains Asian (62%), East African (14%) and West African accessions (11%). Clade 11 contains Asian (57%), Southern African (29%) and West African accessions (14%). Clade 12 contains East African (68%) and Asian accessions (16%). Clade 13 contains primarily East African accessions (83%). Clade 14 contains Southern African (70%) and East African accessions (11%). Clade 15 contains primarily Southern African accessions (80%). Clade 16 contains East African (49%) and West African accessions (30%). Clade 17 contains Asian (35%) American (29%), and West African accessions (17%). Clade 18 contains American (40%) and Asian accessions (39%). Clade 19 contains West African (48%) and Southern African accessions (38%). Clade 20 contains American (66%), Asian (11%) and West African accessions (11%). Clade 21 contains American (59%) and West African accessions (28%). Clade 22 contains West African (30%), American (27%), Asian (15%), and Southern African accessions (13%). Clade 23 contains West African (53%), Southern African (19%), and American accessions (12%). Clade 24 contains American (72%) and Asian accessions (11%). Clade 25 contains Asian (48%) and American accessions (38%). Clade 26 contains American (59%), West African (12%) and Asian accessions (11%). Clade 27 contains West African (62%) and Central African accessions (11%). Clade 28 contains four Asian accessions. Clade 29 contains accessions primarily from Western Africa (84%) with a small representation of Central Africa (11%). Clade 30 contains Asian (27%), American (22%), European (18%) and Southern African accessions (14%). Clade 31 contains only Asian (64%), East African (20%), and West African accessions (16%). Clade 32 contains Asian (70%), North African (10%) and West African accessions (10%). Clades 33 and 34 contain mostly Asian accessions (77% and 92%, respectively; Table S14).

### Improvement status

There is a general trend for Clades 16 – 18, 22 and 23 containing many (>10%) Advanced/Improved Cultivars (12% – 19%), Breeding Lines (11% – 25%), and Landrace/Traditional Cultivars (37% – 68%). Clades 1 – 3, 5 – 8, 11, 20, 24, 26, 27, 30 and 33 contain many (>10%) Advanced/Improved Cultivars (11% – 41%) and Landrace/Traditional Cultivars (26% – 71%). Clades 19 and 21 contain many (>10%) Breeding Lines (19% – 24%) and Landrace/Traditional Cultivars (24% – 62%). Clades 4, 9, 12 – 15 and 32 contain many Landrace/Traditional Cultivars (74% – 100%). Clades 10, 25, 28, 29, 31 and 34 contain many (>10%) Landrace/Traditional Cultivars (43% – 58%) and Unknown (25% – 52%) accessions. Finally, Clade 28 contains four Unknown accessions. Clade 15 contains the most Wild (12%) accessions while Clade 33 contains the most Weedy (3.5%) accessions (Table S15).

### Collection

As the majority of accessions were sourced from USDA (63%), all clades contain a large proportion of accessions from USDA (50% – 100%). However, there were some clades that contained large proportions of accessions from other collections. Clade 23 contained many

(>10%) AGG accessions (20%). Clades 2 – 6, 8 – 14, 17, 21, 27 and 29 – 31 contained many (>10%) IITA accessions (12% – 50%). Clade 34 contained many (>10%) NARO accessions (14%). Clades 16, 18 – 20 and 22 contained many (>10%) AGG accessions (11% – 22%) and IITA accessions (10% – 14%). Clade 32 contained many IITA accessions (20%) and UCR accessions (10%). Clade 15 contained many IIAM (24%) and IITA accessions (20%). Finally, Clade 33 contained many NARO (24%) and AGG accessions (13%) (Table S16).

Cultivar group

Biflora accessions were found at low frequencies in Clades 17, 24, 27, 30, 33 and 34 (0.2% – 3%). Sesquipedalis accessions were found at low frequencies in Clades 10, 15, 17, 23, 30 and 34 (0.1% – 2%) but Clades 32 and 33 contained many sesquipedalis accessions (10% and 27%, respectively; Table S17).

Clade 5

Clade 5 (223 accessions) is characterised by a high proportion of Group 4 accessions (67%), geographically diverse accessions from Asia (36%), Eastern Africa (27%), Western Africa (16%) and Southern Africa (9%), and a combination of Landrace/Traditional Cultivars (67%) and Advanced/Improved Cultivars (20%).

Clade 9

Clade 9 (214 accessions) is similar to Clade 5 and is characterised by geographically diverse accessions from Eastern Africa (36%), Southern Africa (29%), Asia (20%) and Western Africa (11%), and a combination of Landrace/Traditional Cultivars (82%) and Breeding Lines (6%).

Clade 10

Clade 10 (301 accessions) is characterized by a high proportion of Asian (62%), East African (14%) and West African (11%) accessions primarily from India (59%), Kenya (9%) and Nigeria (6%).

Clade 15

Clade 15 contains 894 accessions enriched for Southern African accessions primarily from Botswana and Mozambique (*n* = 288 and 224, respectively) and consists largely of Landrace/Traditional Cultivars (*n* = 660) and Wild accessions (*n* = 108) with fewer Advanced/Improved Cultivars (*n* = 67) and Breeding Lines (*n* = 24). There are 6 sub-clades within clade 15, primarily structed by region of origin (Figure S10). Sub-clade 15a (*n* = 82) contains accessions from a mixture of countries with large representations from Malawi (*n* = 23), Brazil (*n* = 13) and Nigeria (*n* = 10), and contains small clades of Brazilian and Malawian accessions. Sub-clade 15b (*n* = 70) contains accessions primarily from Malawi (*n* = 15) and India (*n* = 10) and but no distinct geographic trends. Most accessions in Sub-clades 15a and 15b are Landrace/Traditional Cultivars (79% combined) with a few Breeding Lines (8% combined). Sub-clade 15c (*n* = 232) contains accessions primarily from Mozambique (*n* = 212) with a small clade of accessions from Zimbabwe. Almost all (99%) of accessions in Cub-clade 15c are Landrace/Traditional Cultivars. Sub-clade 15d (*n* = 185) contains a Botswanan clade (*n* = 39) and multiple small clades from Zambia, Zimbabwe and a combination of the three countries. A combination of Landrace/Traditional Cultivars (72%) and Wild accessions (23%) are present in Sub-clade 15d. Sub-clade 15e (*n* = 137) consists of small clades from Keyna, Lesotho, and Botswana and Lesotho accessions and contains a combination of Landrace/Traditional Cultivars (64%), Advanced/Improved Cultivars (15%),

Wild accessions (9%) and Breeding Lines (5%). Sub-clade 15f ($n$ = 188) consists of two clades, one with accessions from Kenya and Uganda, and the other with primarily Botswanan accessions ($n$ = 133). Similar to Sub-Clade 15e, Sub-clade 15f contains a combination of Landrace/Traditional Cultivars (48%), Advanced/Improved Cultivars (20%) but more Wild accessions (27%). Sub-clade assignment can be found in Table S6 and visualised in Figure S10. All 350 Group 1 accessions and over half of the Wild accessions (58.7%) are found in Clade 15, and the majority of accessions from the IIAM collection are captured in this clade (85%).

Clade 17

Clade 17 contains 1,056 accessions characterised by a high proportion of Group 7 and 8 accessions (16% and 12%, respectively, Table S6), from Asia (35%) the Americas (29%) and Western Africa (17%), and a combination of Landrace/Traditional Cultivars (41%) and breeding material (Advanced/Improved Cultivars and Breeding Lines, collectively 29%) (Figure S11). Clade 17 contains 29% of the accessions from the LAN collection while Clades 20 and 30 contain 37% and 22%, respectively, of LAN accessions.

Clade 20

Clade 20 contains 384 accessions primarily from the Americas (66%), Asia (12%) and Western Africa (12%), and breeding material (Advanced/Improved Cultivars and Breeding Lines, collectively 34%) and Landrace/Traditional Cultivars (26%) (Figure S12). One third ($n$ = 49) of the Group 9 accessions are found in Clade 20 (Table S13).

Clade 24

Clade 24 contains 228 accessions and the majority of Group 9 accessions (66%). Clade 24 accessions are primarily from the USA (63%) and Nigeria (8%) and are Advanced/Improved Cultivars (38%) or Landrace/Traditional Cultivars (28%).

Clades 27 and 29

West and Central African accessions are primarily found in two Clades. Clade 27 (733 accessions) is characterised by a high proportion of Group 5 accessions (63%), originating from Nigeria (28%) and Ghana (13%), and are mainly Landrace/Traditional Cultivars (66%) or Advanced/Improved Cultivars (11%) (Figure S13). Clade 29 is the largest clade, containing 1,874 accessions, primarily from Nigeria (46%), Niger (21%) and Cameroon (8%). All Group 2 accessions ($n$ = 976) are found in Clade 29; 58% of Clade 29 accessions are Landrace/Traditional Cultivars; and 35% of Clade 29 accessions originate from the IITA Genebank (Figure S14).

Clade 30

Clade 30 is the second largest clade containing 1,241 accessions primarily from Group 6 (43%) and contains subclades characterised by geographic origin and contains accessions from: USA (18%), Italy (12%), Iran (11%), Botswana (10%) and Egypt (6%). Clade 30 contains a mixture of Landrace/Traditional Cultivars (48%) and breeding material (Advanced/Improved Cultivars and Breeding Lines, collectively 20%). Clade 30 contains 18% of the accessions from AGG, with Clades 17, 33, and 29 containing 16%, 14% and 11% of AGG accessions respectively (Figure S15).

Clades 33 and 34

Clade 33 contains 712 accessions primarily from Group 3 (60%) and primarily originating from Asia (77%), specifically: India (24%), Japan (15%), China (10%), Philippines (7%), USA

(4%), Taiwan (4%) and Thailand (4%). Ninety-three per cent of the World's classified *Sesquipedalis* (yardlong or asparagus bean) accessions are found in Clade 33, of which, they originate primarily from: China (28%), India (14%), Taiwan (13%) and the Philippines (11%). Fifty-seven per cent of classified *Biflora/Cylindrica* (catjang) accessions are present in Clade 33. Most accessions in Clade 33 are Landrace/Traditional Cultivars (47%) or Advanced/Improved Cultivars (18%). Sixty-eight per cent of the World's Weedy cowpea accessions originate in Clade 33. Half of the accessions in Clade 33 originate from the USDA (50%), with NARO and AGG contributing 24% and 13%, respectively (Figure S16). Similarly, Clade 34 accessions originate from NARO (14%) but primarily originate from the USDA (72%). Clade 34 contains 748 accessions primarily from Group 3 (61%) and originating from Asia (92%), specifically: India (67%), Nepal (12%), Pakistan (5%) and Yemen (3%). Many accessions in Clade 34 are Landrace/Traditional Cultivars (54%) (Figure S17).

Southern African germplasm

Interestingly, many Advanced/Improved Cultivars, Landrace/Traditional Cultivars and Wild accessions originating from Botswana are phylogenetically close (Figure S10), reflecting substantial breeding activity, possibly including introgression from wild cowpea. Africa is particularly rich in wild cowpea diversity (Zuluaga et al., 2021) however studies on the use of wild cowpea relatives in developing improved varieties are scarce, as breeders have avoided them due to undesirable traits such as small seed size, unattractive seed coat colour and texture, disease susceptibility, pod shattering, and indeterminate growth (Rawal et al., 1976). With the development of genomic tools for cowpea, breeders can now more effectively engage in pre-breeding activities to harness the benefits of wild relatives. For example, large seeds are generally preferred and is a common goal for selection. Wild cowpea typically have small seeds but a higher number of seeds per pod. More ovaries per pod is a heritable trait (Drabo et al., 1985) and can be selected for in breeding programs to contribute to higher grain yield. Although small seed size is dominant over larger sizes and controlled primarily by additive gene action with significant epistatic effects (Drabo et al., 1984, Lo et al., 2018), many molecular tools now allow for the elimination of linkage drag, enabling the incorporation of beneficial traits from wild relatives while minimizing undesirable characteristics. Wild relatives can contribute to additional beneficial traits, including climate adaptation in locally adapted germplasm. Macharia et al. (2025) identified Southern African cowpea landraces possess environmental stress tolerant loci (Sub-clade 15c). Therefore, it is possible that the Botswanan wild and landrace accessions in Sub-clades 15e and 15f contain locally adapted loci that have been incorporated into advanced breeding lines.

In contrast, other Southern African countries such as Mozambique contain many landraces but few advanced/improved cultivars or breeding lines (Figure S10). There are limited breeding programs in Mozambique despite it being the second most cultivated legume crop in the country (Gomes et al., 2020). Instead, farmers maintain landraces which usually have high levels of genetic variation because of the many years of uncontrolled cross-regional and infield genetic exchange with past varieties rather than an extended period of selection (Gomes et al., 2020). Substantial genetic variation exists in Mozambique, potentially due to multiple introductions (e.g., accessions spread across clades 9, 15 and 29) and subsequent local adaptation.

Previous studies have hypothesised that cowpea landraces in Portuguese-speaking countries would share a common genetic background (Guimarães et al., 2023). Similar to Guimarães et al. (2023), our phylogenetic tree identified limited evidence suggesting Portuguese and Mozambique landraces share a similar genetic background (Table S6).

Portuguese landraces were more genetically similar to other Mediterranean cowpea landraces (Figure S15), also identified by Carvalho et al. (2017). Additionally, limited evidence in Sub-clades 15a (Figure S10) and 17a (Figure S11, Table S6) suggests similarity between a few accessions from Brazil and Mozambique. Instead, accessions from Brazil and Mozambique were identified in different clades suggesting independent origins and adaptation to local conditions.

### American and Mediterranean germplasm

Genotyping a global cowpea collection has provided insights into its history in the Americas. In South America, accessions primarily originate from five clades ($n > 10$ in Clades 15, 17, 20, 21 and 30) and show regional patterns. As mentioned above, Brazilian accessions originate from multiple origins (Clades 15, 17, 20 and 21), with suggestion for local adaptation occurring (e.g., a unique Brazilian Sub-clade 21b). Argentina, Colombia, Bolivia and Paraguay contain accessions in Clades 17 and 20; while Peru has accessions in Clades 20 and 30. In Central America and the West Indies, accessions primarily originate from five clades ($n > 10$ in Clades 17, 20, 24 and 30). Mexican accessions are present in Clades 17, 20 and 30. In the USA, Breeding Lines from the West (California and Arizona) are primarily in Clade 30 ; Breeding Lines and Landraces from the Midwest (Kansas) are primarily found in Clade 17; and accessions from Southern States are split between Clades 17 (Georgia), 18 (Maryland), 20 (Georgia), 24 (Alabama, Florida, Georgia, Mississippi and South Carolina), 26 (Georgia and South Carolina) and 30 (Alabama, Arkansas, Georgia and South Carolina; Table S18).

This regional structure, along with textual evidence from Herniter et al. (2020), supports the dispersal of cowpea from major gene pools from sub-Saharan Africa to America aligning with historic trade routes. However, this study shows that West African cowpea is largely confined to northern South America, while accessions in most of South America trace their origins to Southeast Africa similar to Carvalho et al. (2017). At least two distinct routes have been proposed for cowpea introduction into the USA: one involving Spanish explorer Hernando de Alcorón in 1540, who introduced it to the Southwest from Mexico (aligning with many Mexican accessions from Clades 17, 20 and 30, Table S18), potentially followed by Jesuit monk Eusebio Kino in the late 1600s (Herniter et al., 2020). The other through the transatlantic slave trade (aligning with many Central and South American accessions in Clades 17, 20, 24, 30 and 33, Table S18), bringing it as provisions to the Southeastern United States (Herniter et al., 2020). This dual mode of introduction, is supported by the findings of Muñoz-Amatriaín et al. (2021) and Herniter et al. (2020). However, in this study we identified that American accessions are derived from at least four major lineages (Clades 17 ($n = 303$), 30 ($n = 269$), 20 ($n = 254$) and 24 ($n = 164$)) reflecting both historic and recent introductions. Genetic distinctions between these multiple introductions into the Americas are evident, aligning with Carvalho et al. (2017), which identified genetic similarities between African and South American cowpea varieties, suggesting multiple routes of dispersal during the colonial era.

### Asian germplasm

The earliest evidence of cowpea outside of Africa dates to 3,500 BP in Daimabad, India (Fuller, 2003), and textual evidence indicates its presence in the Mediterranean Basin by at least 2,500 BP, during the time of the Ancient Greeks (Herniter et al., 2020). Cowpea's spread into India is proposed to have occurred via two potential routes: (1) the Sabaean lane, which originated in Africa, passed through modern Yemen, and utilised monsoonal winds to cross the Arabian Sea, or (2) through inland trade routes, moving from Egypt to the Near East and then across the Iranian Plateau to northwest India. Archaeological and linguistic evidence suggests that the Sabaean route is more likely, indicating that Asian cowpea would be more closely related to West and North African cowpea. Genetic evidence from this study confirms cowpea in Asia are closely related to West African cowpea (Figure 4). However, some Asian (Iran, Pakistan and Afghanistan) and Mediterranean cowpea are closely related (Figure S15), suggesting that after cowpea reached the Mediterranean region, subsequent re-introduction into Asia occurred possibly along trade routes such as the Silk Road. Multiple trade routes both historic and recent, convolute the relationships among extant cowpea subsequently obfuscating climatic adaptation and local selection.

As cultivated cowpea spread into Asia, selection for fresh pod consumption led to the divergence of the “yard-long” cultivar group (*V. unguiculata* ssp. *unguiculata* cv. *sesquipedalis*). Most *sesquipedalis* accessions (78%) were grouped into Group 3 at the $K$ = 9 level, with the remaining 22% admixed among other groups. The majority of these accessions were from Eastern Asia, consistent with the geographic distribution of the cultivar (Table 1). The clear separation of the *sesquipedalis* cultivar group supports the hypothesis that it diverged from grain cowpea due to selection for traits favourable to vegetable use, such as long tender pods. The spread of cowpea in Asia occurred primarily from India to North (Nepal and Pakistan; Figure S17), East and Southeast Asia (Figure S16), with vegetable cowpea then dispersing from China to the rest of Asia and America (Figure S16). Geographic clustering suggests that regional breeding practices and environmental factors have significantly influenced genetic variation, emphasizing the need for future analyses to account for geographic structure, particularly in studies such as GWAS. These findings highlight the significant role of East and Southeast Asia in the transmission and diversification of cowpea across the region.
